# Pre-existing partner-drug resistance facilitates the emergence and spread of artemisinin resistance: a consensus modelling study

**DOI:** 10.1101/2021.04.08.437876

**Authors:** Oliver J Watson, Bo Gao, Tran Dang Nguyen, Thu Nguyen-Anh Tran, Melissa A Penny, David L Smith, Lucy Okell, Ricardo Aguas, Maciej F Boni

**Affiliations:** MRC Centre for Global Infectious Disease Analysis, Faculty of Medicine, Imperial College, London; Department of Pathology and Laboratory Medicine, Brown University, Providence, RI, USA; Centre for Tropical Medicine and Global Health, Nuffield Department of Medicine, University of Oxford, Oxford, UK; Center for Infectious Disease Dynamics, Department of Biology, Pennsylvania State University, University Park, PA, USA; Swiss Tropical Public Health Institute, Basel, Switzerland; Department of Health Metrics Sciences, University of Washington, Seattle, WA, USA

**Keywords:** malaria, drug resistance, artemisinin-combination therapies, individual-based modeling, consensus exercise

## Abstract

**Background:** Artemisinin-resistant genotypes have now emerged a minimum of five times on three continents despite recommendations that all artemisinins be deployed as artemisinin combination therapies (ACTs). Widespread resistance to the non-artemisinin partner drugs in ACTs has the potential to limit the clinical and resistance benefits provided by combination therapy.

**Methods:** Using a consensus modelling approach with three individual-based mathematical models of *Plasmodium falciparum* transmission, we evaluate the effects of pre-existing partner-drug resistance and ACT deployment on artemisinin resistance evolution. We evaluate settings where dihydroartemisinin-piperaquine (DHA-PPQ), artesunate-amodiaquine (ASAQ), or artemether-lumefantrine (AL) are deployed as first-line therapy. We use time until 0.25 artemisinin resistance allele frequency (the establishment time) as the primary outcome measure.

**Findings:** Higher frequencies of pre-existing partner-drug resistant genotypes lead to earlier establishment of artemisinin resistance. Across all scenarios and pre-existing frequencies of partner-drug resistance explored, a 0.10 increase in partner-drug resistance frequency on average corresponded to 0.7 to 5.0 years loss of artemisinin efficacy. However, the majority of reductions in time to artemisinin establishment were observed after the first increment from 0.0 to 0.10 partner-drug resistance genotype frequency.

**Interpretation:** Partner-drug resistance in ACTs facilitates the early emergence of artemisinin resistance and is a major public health concern. Higher grade partner-drug resistance has the largest effect, with piperaquine-resistance accelerating early emergence of artemisinin-resistant alleles the most. Continued investment in molecular surveillance of partner-drug resistant genotypes to guide choice of first-line ACT is paramount.

**Funding:** Bill and Melinda Gates Foundation; Wellcome Trust.

## Introduction

Worldwide adoption of artemisinin-combination therapies (ACTs) against uncomplicated *Plasmodium falciparum* malaria began after the World Health Organization (WHO) recommended ACTs as the first-line therapy in all malaria-endemic countries in 2005 ^1^. Prior to 2005 artemisinin was being primarily used in Southeast Asia – both as a monotherapy and in combination therapies. The emergence of artemisinin resistance in the Greater Mekong Subregion (GMS) has been largely attributed to this longer history and higher frequency of use of artemisinin-derivatives compared to Africa, particularly as monotherapy. As an example, despite the national switch to artesunate-mefloquine (ASMQ) in 2000, one report revealed that 78% of all artemisinin delivered in Cambodia in 2002 was as a monotherapy ^2^. This uncontrolled artemisinin monotherapy use in private market drug purchases increased the risk of artemisinin resistance emerging, with no partner drug to protect artemisinins should an artemisinin-resistant genotype emerge. Despite efforts to remove monotherapies from private markets and clinical use ^3^, the first documented cases of artemisinin resistance were observed in western Cambodia in 2008 ^4^, although the resistant genotype had emerged years earlier and was already at high frequency in 2002 ^5^. Currently, the majority of artemisinin-resistant genotypes are confined to SE Asia, however, independent emergence of artemisinin resistance has now been identified in both Guyana ^6^ and New Guinea ^7^. The first observation of de novo emergence of *pfkelch13*-mediated artemisinin resistance in Africa was made in Rwanda in 2020 ^8^.

In response, the vast majority of artemisinin is administered in the form of ACTs. A key role of the non-artemisinin partner drug is to reduce parasite densities of emergent artemisinin-resistant genotypes ^9^. However, in epidemiological scenarios where *P. falciparum* is resistant to these partner drugs ^10^, ACTs may already be acting as de facto monotherapies 11. Mutations conferring high level resistance to the partner drugs piperaquine and mefloquine have already spread in some areas of the GMS. In Africa, the most commonly used partner drugs are lumefantrine and amodiaquine, for which partial resistance has been observed in multiple settings ^12^. High levels of ACT use in areas with high frequencies of partner-drug resistance may (1) pose an increased risk of artemisinin-resistance *de novo* emergence and subsequent spread, and (2) create conditions where imported artemisinin-resistant genotypes may rapidly spread leading to elevated treatment failure. By contrast, choosing ACT policy according to local partner drug efficacy may reduce resistance spread. Here, we evaluate the long-term effects of partner-drug resistance evolution on selection pressure and early emergence of artemisinin-resistant genotypes. We use a consensus approach taken in previous mathematical modeling studies ^13,14^ and present results from three independently-built individual-based simulations of *P. falciparum* malaria.

## Methods

### Model descriptions

Three individual-based stochastic models of malaria transmission and evolution were used to evaluate artemisinin-resistance evolution under different pre-existing levels of partner-drug resistance. The three models were developed by the MRC Centre for Global Infectious Disease Analysis, Imperial College London (‘Imperial’ ^15^), the Center for Infectious Disease Dynamics (CIDD) at Pennsylvania State University (‘PSU’ ^11^), and the Mahidol-Oxford Research Unit (MORU) affiliated with the University of Oxford (‘MORU’ ^16^). Each model simulates 100,000 individuals with a daily time-step that updates individuals’ infection status, treatment status, immunity, genotype-specific parasite densities, and clinical state. For each scenario, models are run until a steady-state prevalence is achieved prior to evaluating resistance evolution over a 40-year period.

Each model tracks the dynamics of clonal blood-stage *P. falciparum* populations within individuals. A key common feature is that individuals can be infected with multiple clones simultaneously. Each parasite clone is described by a genotype that characterises their antimalarial resistance phenotype. These include the K76T locus in *pfcrt*, N86Y and Y184F in *pfmdr1*, C580Y in *pfkelch13*, copy number variation (CNV) of the *pfmdr1* gene, and CNV of the *pfpm2-3* gene(s). CNV is treated as a binary variable, distinguishing between single-copy and multiple-copies, resulting in 64 possible genotypes that are tracked. Models allow for treatment of febrile malaria with three different ACTs: artemether-lumefantrine (AL), artesunate-amodiaquine (ASAQ), and dihydroartemisinin-piperaquine (DHA-PPQ). The same parameterization in ^17^ of 3 × 64 = 192 treatment efficacies of three therapies on 64 genotypes was used in all three models. Pleiotropy at the *pfcrt* and *pfmdr1* loci ^12,18,19^ was accounted for with the effects of *pfcrt* and *pfmdr* genotypes on efficacy of both lumefantrine and amodiaquine explicitly modelled. The C580Y locus is used as a proxy for artemisinin resistance, recognizing that multiple *pfkelch13* mutations have been associated with a drop in artemisinin efficacy. Full model details can be found in Supplementary Materials.

### Alignment

We carried out an alignment exercise to ensure model outputs were comparable across the explored epidemiological scenarios. We aligned the three models’ de novo mutation rates so the models reached 0.01 580Y allele frequency (early artemisinin resistance emergence (Supplementary Figure 1)) after seven years exactly, under a specified set of conditions: 100,000 individuals in a transmission setting with all-ages PfPR=10% and 40% coverage with DHA-PPQ as a first-line therapy (Supplementary Figure 2). This was chosen as the *de novo P. falciparum* mutation rate and within-host competition of new mutants cannot be measured precisely enough to simply set each model’s mutation and fixation parameters to a particular value. This alignment also ensures that one model does not generate a larger number of drug-resistant mutants simply because its mutation rate is higher. We also demonstrate that the 0.01 allele frequency threshold is insensitive to selection pressure for the mutation rates explored (Supplementary Figure 3), validating it as an appropriate cross-model alignment of the combined process of de novo mutation and within-host fixation of new alleles. In addition, by using a common drug-by-genotype efficacy table ^17^ we ensure that the post-emergence treatment failure patterns are similar across all models (Supplementary Methods and Supplementary Table 1). Treatment failure is defined as PCR-corrected recrudescence after 28-days, as recommended by the WHO for surveillance of antimalarial efficacy ^20^. Crucially, by conducting our alignment this way, we followed similar model consensus exercise methodology that assures the compared models are not aligned on their defined outcome measures, and instead focus on evaluating the comparative impact of different epidemiological scenarios on each model ^13,14^.

**Table 1.** Times till 0.25 580Y frequency (T_Y0.25_) under 40% treatment coverage under different starting partner drug (PD) resistance. Mean years for scenarios with 0 starting partner drug resistance (PD_0_) is given before showing the mean percentage difference in T_Y0.25_ for each partner drug resistance explored. Range in square brackets reflects the interquartile range across all prevalence settings considered. Censored times above 40 years were inferred using a Weibull distribution to describe time-to-event values.

| Model | ACT | Years till 0.25 580Y frequency | Percent reduction in establishment time (or artemisinin UTL) when comparing to $PD_0$ scenario | | | | Years Lost per 0.10 PD increase |
| --- | --- | --- | --- | --- | --- | --- | --- |
| | | $T_{Y0.25}, PD_0$ | %$T_{Y0.25}, PD_{0.01}$ | %$T_{Y0.25}, PD_{0.1}$ | %$T_{Y0.25}, PD_{0.25}$ | %$T_{Y0.25}, PD_{0.5}$ | |
| MORU | DHA-PPQ | 25.3 [21.6, 33.2] | 24.3% [17.7, 30.8] | 50.4% [44.9, 56.4] | 58.8% [52.7, 64.0] | 63.4% [57.5, 69.2] | 3.2 |
| PSU | DHA-PPQ | 13.9 [11.8, 17.1] | 2.4% [-1.6, 6.5] | 15.4% [11.8, 19.0] | 20.3% [16.5, 23.7] | 24.6% [21.1, 28.0] | 0.7 |
| Imperial | DHA-PPQ | 27.7 [25.5, 30.2] | 8.4% [5.2, 11.7] | 25.0% [22.2, 27.7] | 30.9% [28.4, 33.8] | 34.1% [31.6, 36.7] | 1.7 |
| MORU | ASAQ | 35.4 [26.7, 51.2] | 12.0% [5.1, 19.0] | 48.1% [42.9, 53.3] | 59.1% [53.8, 64.2] | 66.5% [61.4, 71.5] | 5.0 |
| PSU | ASAQ | 39.4 [33.1, 45.0] | 1.6% [-1.6, 5.0] | 16.7% [13.6, 20.0] | 28.8% [25.8, 32.1] | 31.6% [28.6, 34.8] | 2.5 |
| Imperial | ASAQ | 36.5 [33.1, 41.5] | 2.7% [-0.9, 6.6] | 4.8% [1.7, 8.3] | 10.2% [6.7, 13.5] | 16.4% [13.2, 19.5] | 1.2 |
| MORU | AL | 28.4 [23.3, 34.6] | 6.3% [0.9, 11.5] | 35.2% [31.3, 39.2] | 46.0% [41.9, 49.4] | 53.5% [50.0, 57.1] | 3.1 |
| PSU | AL | 26.4 [22.3, 32.2] | 0.9% [-3.2, 4.9] | 14.0% [10.0, 17.7] | 21.5% [17.7, 25.2] | 29.3% [25.6, 32.8] | 1.6 |
| Imperial | AL | 27.3 [24.9, 31.5] | 5.3% [1.8, 8.5] | 8.3% [4.8, 11.8] | 13.0% [9.6, 16.4] | 17.8% [14.6, 21.2] | 0.9 |

**Figure 1.**
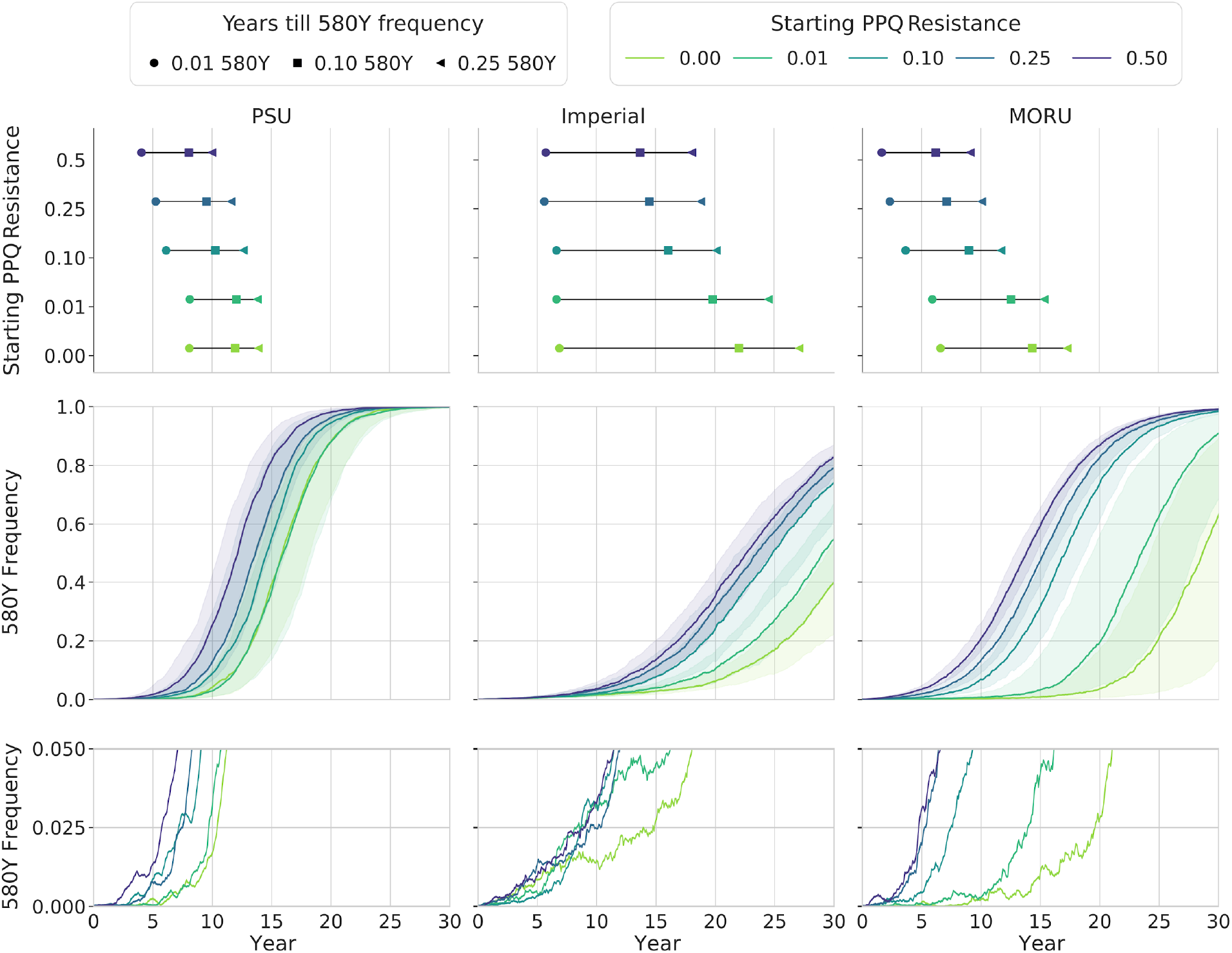
Features of artemisinin selection with respect to starting partner drug resistance frequency. In the scenarios above, DHA-PPQ is used as the first-line therapy, with 40% population level drug coverage and 5% PfPR. Results are shown for five different starting PPQ resistance frequencies (0.00 (green) - 0.50 (purple)). The top row shows the median time to three resistance milestones for the five partner-drug resistance scenarios. The circle, square and triangle show the 0.01, 0.10 and 0.25 artemisinin-resistance frequency milestone respectively. The middle row shows the fixation pattern of artemisinin-resistant genotypes, characterised by the 580Y allele, with the median and interquartile range shown with shaded bands. The bottom row shows the early patterns of emergence for five median simulations, where the’median simulation’ is defined as the one whose time to 0.10 580Y allele frequency is the median time.

**Figure 2:**
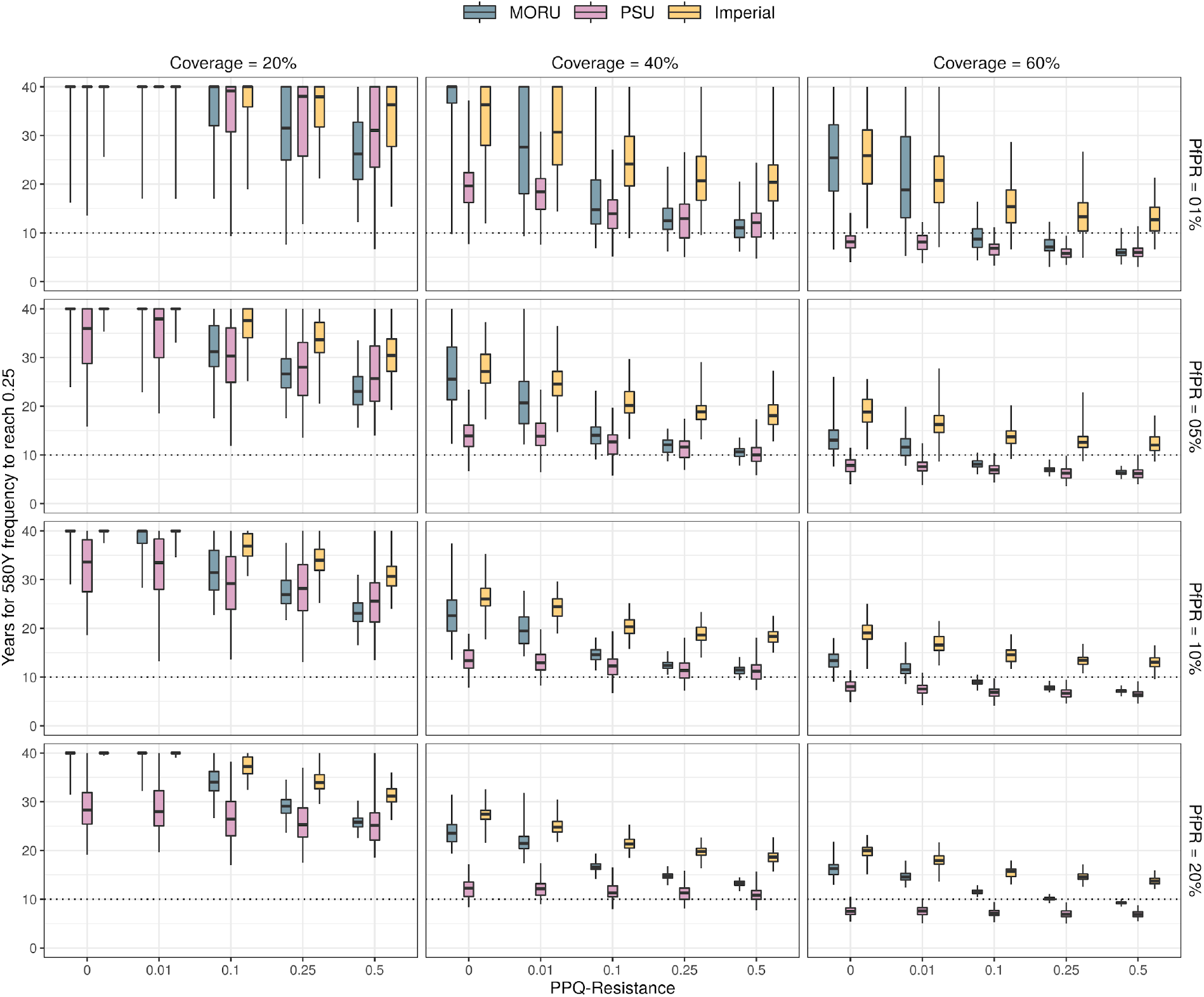
Number of years until 580Y allele frequency reaches 0.25 in regions with DHA-PPQ deployed as first-line therapy, starting from 0.0 580Y frequency. Results for different coverage levels (three columns) and different prevalence (PfPR) levels (four rows) are shown in the 12 panels. The x-axis in each panel shows the pre-existing genotype frequency of plasmepsin CNVs which confer PPQ resistance, and the box plots show the median and interquartile range for the number of years to each 0.25 allele frequency of 580Y, from three different mathematical models. Box-plot whiskers show 95% ranges from 100 simulations. As the initial genotype frequency of PPQ-resistance increases, the time to 580Y establishment gets shorter.

**Figure 3.**
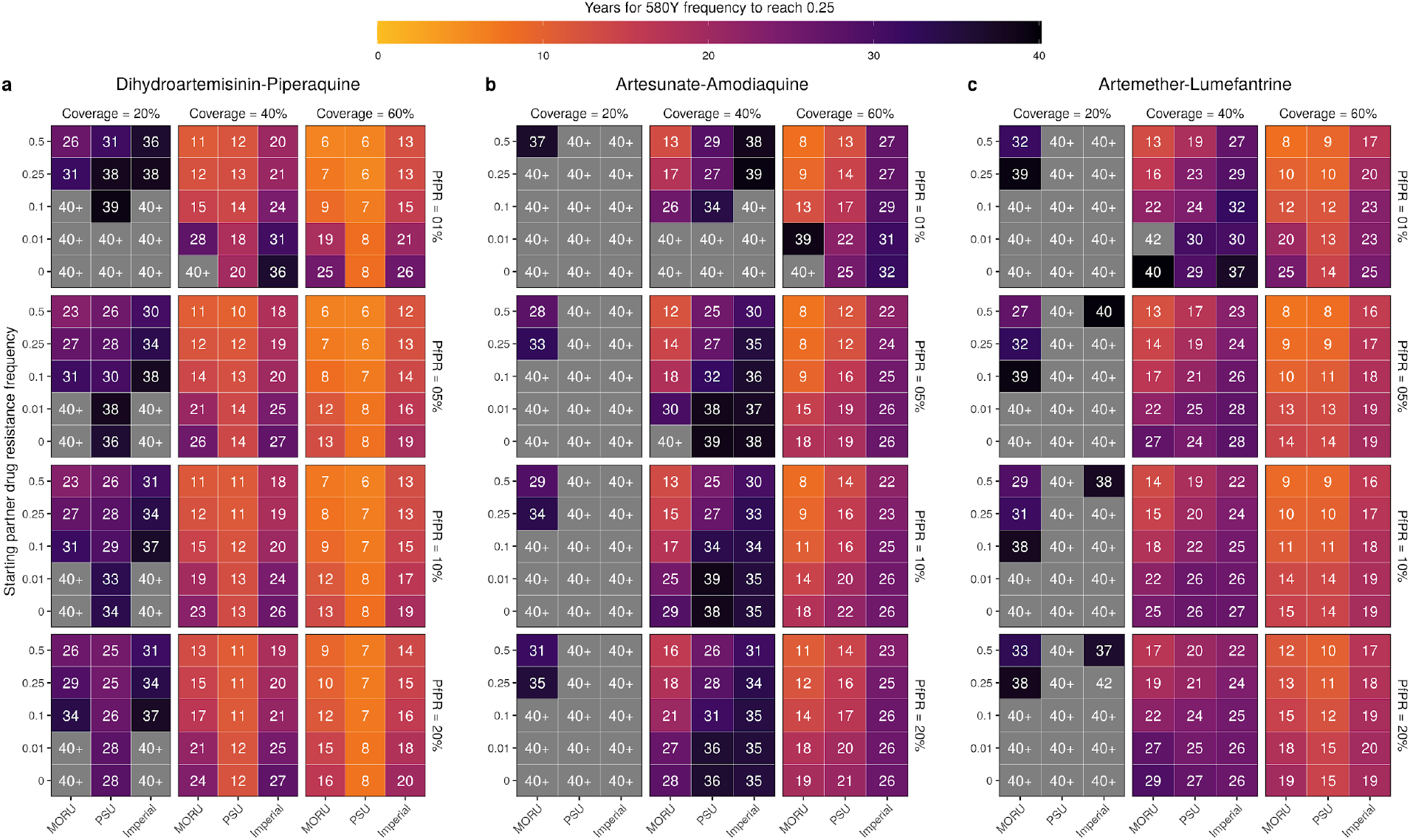
Median time (*T*_*Y*,0.25_) for artemisinin-resistant genotypes to reach a frequency of 0.25, under different conditions of pre-existing partner-drug resistance frequency, starting from 0.0 580Y frequency. The results are shown for three different ACTs; a) DHA-PPQ, b) ASAQ and c) AL. Simulations were evaluated over a 40 year period with median times taking longer than 40 years indicated in grey (40+). Times are shown for each model at four different malaria prevalences (1%, 5%, 10%, 20%) and three different treatment coverages (20%, 40%, 60%). In all settings and across all models a decrease in time to 0.25 artemisinin-resistance frequency was observed with increasing initial partner drug resistance.

### Scenario evaluations

Each epidemiological scenario consists of 100,000 individuals in a particular transmission setting (PfPR=1%, 5%, 10%, 20%), with varying access to antimalarial drugs if febrile (coverage=20%, 40%, 60%). One primary ACT is used as first-line therapy (DHA-PPQ, ASAQ, AL) and five different frequencies of pre-existing partner drug resistance (0.0, 0.01, 0.10, 0.25, 0.50) at time = 0 are explored, with partner drug resistance allowed to spread in response to selection. For piperaquine, pre-existing resistance is defined by the frequency of genotypes with multiple copies of the plasmepsin genes. For amodiaquine, pre-existing resistance is defined by the frequency of genotypes with *pfcrt* 76T, *pfmdr1* 86Y and Y184, which is the most amodiaquine-resistant genotype in our parameterization. For lumefantrine, pre-existing resistance is defined by the frequency of genotypes with *pfcrt* K76, *pfmdr1* N86 184F and double-copy *pfmdr1* genotypes, which is the most resistant lumefantrine-resistant genotype in our parameterization.

### Primary outcome measures

The primary outcome measures were time to reach a particular resistance milestone: time to 0.10 (*T*_*Y*,0.10_) and 0.25 (*T*_*Y*,0.25_) frequency of the 580Y allele (artemisinin resistance indicator). We refer to *T*_*Y*,0.25_ as the establishment time or the useful therapeutic life of artemisinins. Allele frequency is computed in a weighted manner, factoring in monoclonal and multiclonal infections (Supplementary Methods). We also report selection coefficients for the 580Y allele, defined via a sigmoidal fixation curve as follows ^21^:

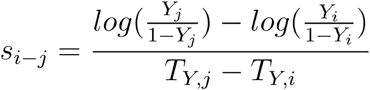

where *Y*_*i*_ and *Y*_*j*_ are the frequency of the 580Y allele at different time points. Here, we present *S*_0.01−0.1_ and *S*_0.1−0.25_ to describe the strength of selection over the time period between reaching 0.01 and 0.10 frequency and between reaching 0.10 and 0.25 frequency. These two periods of evaluation are chosen to show the early period of evolution from low frequencies (0.01) to moderate frequencies (0.10) where random extinction is no longer possible; and the subsequent period to establishment (0.25) after which 580Y alleles are likely to undergo rapid selection and fixation (Figure S1).

Lastly, comparisons of key model assumptions were explored in a sensitivity analysis, with the following assumptions varied: (1) fitness costs associated with resistance, (2) genotype-specific drug efficacies, (3) duration of the asymptomatic infectious period, and (4) probability of developing clinical symptoms after an infection. These facets were chosen as they explore different components within the transmission models at which the selective advantage of drug resistance manifests. Full details described in Supplementary Materials.

## Results

Under long-term deployment of artemisinin-combination therapies in malaria-endemic countries, artemisinin resistance emerges earlier if partner drugs are allowed to fail. An illustration of this process can be seen in Figure 1, where a scenario of 40% treatment coverage and 5% malaria prevalence (PfPR) results in evolutionary paths of artemisinin resistance that emerge and fix earlier under higher levels of pre-existing partner-drug resistance. In these modeled scenarios, the recommended first-line ACT is DHA-PPQ, and only two genetic resistance mechanisms are tracked, the *pfkelch13* 580 locus and copy number of the plasmepsin-2,3 genes. The average rate of selection is approximately the same under different levels of pre-existing piperaquine resistance, since the relative fitness advantage of 580Y over C580 stays approximately constant as C580 alleles are replaced (Supplementary Figure 4). However, with pre-existing piperaquine-resistance at higher levels, the early stochastic stages of artemisinin-resistance emergence are less susceptible to random extinction; this is due to an emergent 580Y allele having higher survivorship when it arrives on a genetic background coding for piperaquine resistance than if emerging alone on a background of partner-drug sensitivity (bottom panels, Figure 1). This means that with higher frequencies of partner-drug resistance, 580Y alleles can establish ‘escape velocity’ (increase in resistance frequency is linear and the probability of extinction is unlikely) earlier and replace drug-sensitive alleles more quickly. This difference between early-phase selection and late-phase selection can be observed by contrasting the approximately flat selection coefficients, with respect to pre-existing piperaquine-resistance, when calculated during the latter phase of selection (S_0.1,−0.25_ in Supplementary Figure 4b) against the sloped selection coefficients observed when calculated during an earlier phase of selection (S_0.01−0.1_ in Supplementary Figure 4a).

**Figure 4.**
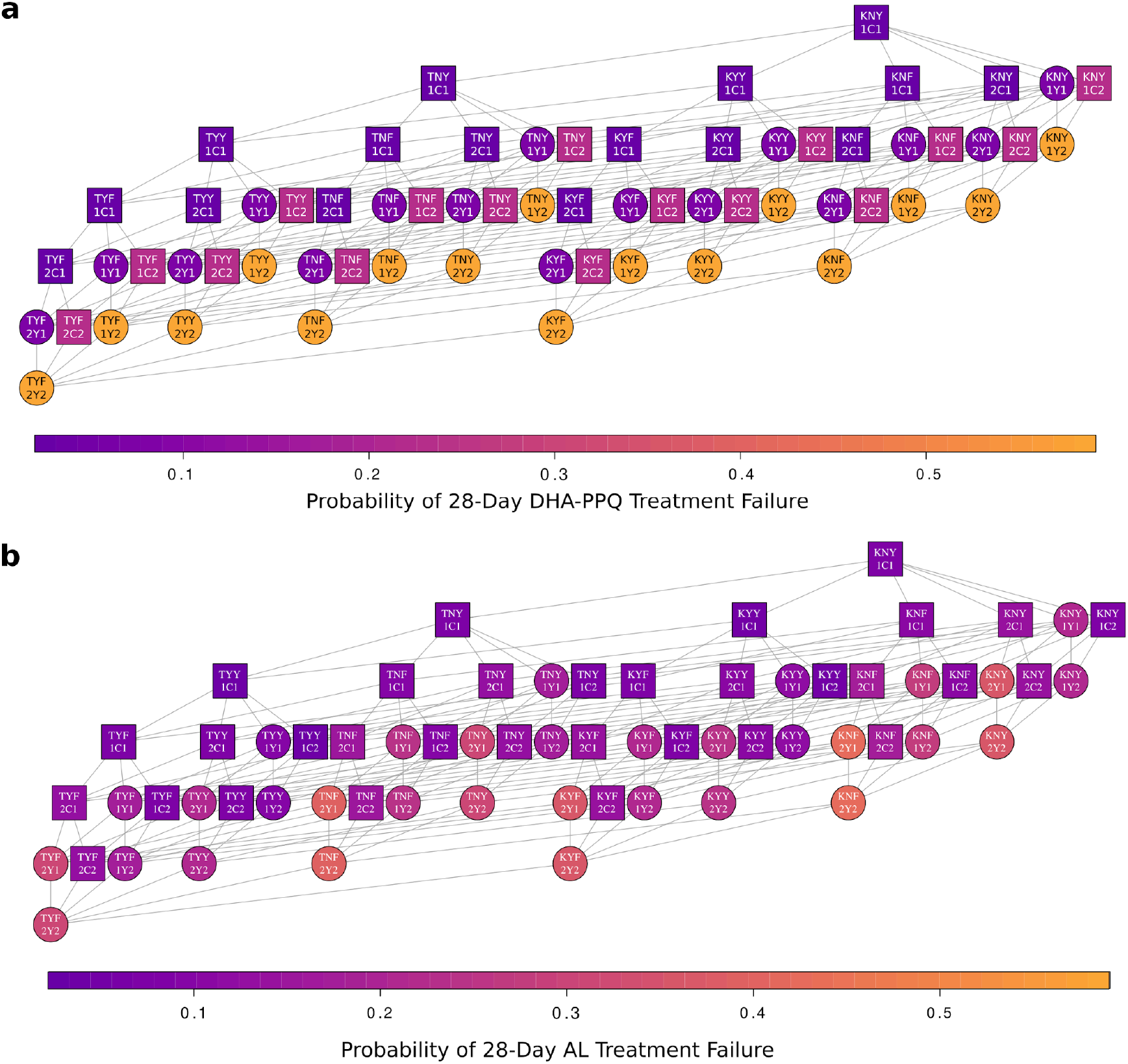
Comparison of fitness landscape of DHA-PPQ and AL. Network diagrams connect genotypes that are one mutation apart and arrange all genotypes from the wild-type (top) to multi-allelic resistant types (bottom); each horizontal level corresponds to one additional genetic mechanism (mutation or CNV) of resistance. The genotype is labelled for each node, e.g. the wild-type at the top, KNY1C1, contains *pfcrt* **K**76, *pfmdr1* **N**86 and **Y**184, **1** copy of *pfmdr1*, *pfkelch* **C**580 and **1** copy of the *pfpm2-3* gene(s). The 28-day treatment-failure probability of each parasite genotype is indicated by the colour of the nodes in the network for a) DHA-PPQ and b) AL, which is detailed in the color bar. Yellow shades show increased resistance (increased probability of treatment failure). Genotypes with the 580Y artemisinin-resistant allele are shown with circles, and genotypes with the wild-type C580 allele are shown with squares. The network highlights the comparatively more complex fitness landscape associated with AL resistance (16 different treatment failure phenotypes) than with DHA-PPQ resistance (4 different phenotypes).

Across transmission and coverage settings, all three models predicted quicker progression to higher frequencies of artemisinin resistance when PPQ-resistance was already high. Under 40% ACT coverage with no pre-existing PPQ-resistance, the median time to establishment (580Y frequency = 0.25) across all four PfPR settings was 26.0 to 36.3 years after introducing DHA-PPQ as first line treatment (Imperial model), 12.3 to 19.6 years after introduction (PSU model), and 22.6 to 40+ years after introduction (MORU model); see Figures 2, 3 and Supplementary Table 3. This duration includes the seven years taken for 580Y alleles to reach 0.01 frequency (Supplementary Figure 2). The variation in durations is a feature of each model’s construction, each using different implementations of the epidemiological, entomological, and clinical aspects of malaria (Supplementary Table 1). When PPQ-resistance is present at 0.10 allele frequency, Figure 2 shows that 580Y establishment occurs 21.9% to 33.6% earlier than under no pre-existing PPQ resistance (Imperial model), 8.1% to 29.1% earlier in the PSU model, and 29.4% to 63.0% earlier in the MORU model. The consistent reduction in establishment time across all three models and four prevalence settings indicates that this is likely a robust feature of *P. falciparum* evolutionary dynamics in modern treatment contexts, namely, that early emergence and establishment of artemisinin resistance is facilitated by the presence of piperaquine resistance in settings where DHA-PPQ is used as first-line therapy.

Rates of artemisinin-resistance evolution differ among the three modeled ACTs (Figure 3). The difference in selection patterns can be explained by the differential efficacy of each ACT as resistance mutations accumulate to the partner drugs, as well as the different number of genetic mechanisms required to achieve maximum levels of resistance (Figure 4) (Supplementary Figure 13). On average, DHA-PPQ treatment efficacy (28-day parasite clearance) drops from 97% (WT) to 93% (580Y) or 77% (plasmepsin CNV) after one resistance mechanism is acquired, and to 42% when both resistance mechanisms are present. These drops correspond to large fitness differences among the parasite genotypes. ASAQ efficacy drops from 98% (WT) to 95% (580Y) or 89% (three AQ resistance mutations), and then to 74% on a double-resistant genotype. AL efficacy drops from 97% to 91% (580Y) or 83% (four LUM resistance mechanisms), and then to 57% efficacy on the double-resistant genotype. Hence, for all three models, resistance evolution under DHA-PPQ occurs the most quickly yielding the greatest selection coefficients (Supplementary Figure 5) as the relative fitness advantages of the resistant genotypes are the highest and the number of mutations required is the fewest. With no pre-existing partner-drug resistance and 40% coverage, median establishment time across all models and prevalence settings was 23.8 years (interquartile range (IQR): 16.2 - 28.3) for DHA-PPQ, 37.1 years (IQR: 31.5 - 44.5) for ASAQ, and 27.3 years (IQR: 23.8 - 32.5) for AL (Table 1, Figure 3).

Time until establishment of artemisinin resistance varied with first-line therapy, malaria prevalence, and treatment coverage (Figure 2). Evolution is always faster under higher treatment coverage, in all models and scenarios (Figure 3). In our modeled scenario runs, resistance evolution is generally but not always faster at higher prevalence levels (Figure 3, Supplementary Figure 6). This observation is most pronounced in scenarios explored at 1% PfPR, which exhibited higher variance in emergence times leading to (on average) longer establishment times for 580Y. In the present analysis, the general trend of resistance evolution occurring more quickly at higher prevalence levels results from more *de novo* mutation and shorter parasite generation times. Establishment times for 580Y show a clear pattern of occurring earlier under higher partner-drug resistance (Table 1). An increase from 0.0 to 0.10 starting partner-drug resistance frequency is associated with a range of 2 to 16 years of lost artemisinin efficacy. Ranges are taken across all prevalence levels, first-line therapies, and models. The number of years of lost artemisinin efficacy is approximately log-linear with respect to increases in partner drug resistance (Supplementary Figure 6). Consequently, the majority of reductions in time to artemisinin establishment are observed after the first 0.10 genotype-frequency increment in pre-existing partner-drug resistance. This observation was true for all scenarios explored except when (i) using AL as first line-therapy in the PSU and Imperial College models, and (ii) using ASAQ in the Imperial College model. For example, in Table 1, only a 14.0% and 8.3% median reduction in establishment time was observed with 0.1 frequency of pre-existing lumefantrine resistance for the PSU and Imperial College model respectively. This represented just 46.6% and 47.8% of the total reduction observed when 0.5 frequency of pre-existing lumefantrine resistance was assumed.

As several model assumptions are known to affect evolutionary dynamics but are difficult to validate or estimate with field data, we conducted sensitivity analyses on the resistant genotypes’ fitness costs, the genotype-specific drug efficacies, the duration of infection of asymptomatic carriage, and the probability of progressing to symptoms after an infectious bite (Supplementary Figures 7 to 10). Although evolutionary rates are affected by these changes -- e.g. higher fitness costs and lower probability of symptoms lead to slower drug-resistance evolution -- the relationship between higher pre-existing partner-drug resistance and earlier 580Y establishment was shown to be robust under all scenarios examined.

## Discussion

Drug-resistance surveillance efforts have often focused on understanding the emergence, spatial spread, and evolution of artemisinin-resistant genotypes. After the discovery of molecular markers underpinning artemisinin resistance in SE Asia ^5^, a number of studies were undertaken to investigate the prevalence of these markers in Africa. In contrast, resistance to partner drugs has received comparatively less attention, despite ACTs being used in epidemiological settings with endemic resistance to amodiaquine, lumefantrine, and mefloquine. Our results show that focussing surveillance on partner-drug resistance is needed to prevent the spread of artemisinin resistance.

Here, we show that emergence and spread of partner-drug resistance gradually erodes both ends of an ACT’s useful therapeutic life - via immediate reductions in efficacy due to lower partner-drug pharmacodynamic activity and by shortening the period when ACTs can be used at full efficacy before artemisinin-resistant genotypes are established. Using three independently-calibrated stochastic individual-based malaria models, the primary evolutionary effect appears to occur at low 580Y allele frequencies. During this early emergence period, artemisinin-resistant genotypes are able to establish more quickly when appearing on a genetic background of partner-drug resistance, but have delayed establishment when appearing alone and experiencing relatively high ACT treatment efficacies (91% to 95%). An analysis of selection coefficients during the establishment phase (580Y frequencies > 0.10) also shows an additional but smaller effect of stronger selection of 580Y alleles when appearing alongside partner-drug resistance mutations (Supplementary Figure 4a).

Substantial reductions in the establishment time for artemisinin resistance was observed after a 0.10 increase in partner-drug resistance frequency. This finding was broadly similar across models and first-line scenarios, with the majority of reductions in time to artemisinin establishment observed with only 0.10 pre-existing partner-drug resistance. These effects are of most practical consequence for DHA-PPQ, as evolution of artemisinin-resistance is fastest (in all three models) in scenarios where DHA-PPQ is deployed as first-line therapy. Copy-number variation at the plasmepsin loci corresponds to larger drops in ACT efficacy ^10^ than those observed for lumefantrine-resistant and amodiaquine-resistant genotypes ^17^. A second reason for the earlier emergence of genotypes fully-resistant to DHA-PPQ is that in all three models they require only one mutation and one CNV, whereas complete resistance to ASAQ and AL requires three mutations, with smaller fitness differences associated with each individual mutational step.

The current distribution of partner-drug resistance is not well characterised in all countries and is dynamic given collateral sensitivity between drugs. The partially resistant amodiaquine haplotype is common throughout South-East Asia and Africa, in part driven by historical chloroquine use ^22^. However, there is some evidence of its decline in recent years particularly since widespread use of AL in Africa which selects in the opposite direction at these loci ^12^. Multi-copy *pfmdr1* parasites were originally more common in South-East Asia but appeared to decline when artesunate-mefloquine was first-line treatment; they are now increasingly being observed in Africa ^23^. Piperaquine resistance reached high levels in Cambodia and bordering areas. Surprisingly high levels of multiple-copy plasmepsin have also been observed in several countries in Africa, despite apparent high efficacy of DHA-piperaquine in the same areas ^24^. More surveillance and efficacy studies are required to both reconcile these results and map all relevant resistance markers.

Additional evolutionary mechanisms are likely to play a role in navigating the multi-drug resistance pathway to ACT resistance (Figure 4). Clonal competition is likely to slow down the emergence of drug resistance, with new genotypes competing within-host in multiclonal infections and between-host to rise to high frequencies in the population (Supplementary Figure 10). This is particularly important if resistance incurs a fitness cost, meaning newly emerging drug resistant parasites may be outcompeted by fitter drug sensitive clones when drug concentration is below inhibitory concentration ^25^. Second, the likelihood of mutation events occurring in the absence of drug pressure and being onwardly transmitted will affect the generation of additional partner-drug resistant genotypes and impact the speed of selection due to increased interclonal interference in the population (Supplementary Figure 12). Third, genetic recombination can act to both unite and break apart multi-genic resistant genotypes. Population genetics approaches are divided on whether recombination would speed up or slow down the arrival of multi-drug resistant genotypes ^26,27^. An improved understanding of these mechanisms will help us gain an understanding as to which field observations -- e.g. step-wise mutation patterns observed in high transmission regions -- present the most or least risk for drug resistance emergence at different transmission intensities. Our modeled patterns of resistance evolution did not show a consistent relationship with transmission intensity, suggesting that monitoring efforts should be supported wherever possible. Although low-transmission regions have historically been associated with drug-resistance emergence ^28^, our findings are consistent with both empirical and theoretical work suggesting a non-monotonic relationship between malaria prevalence and resistance evolution ^29,30^.

Our structured model comparisons have several important limitations. First, drug-resistant genotypes (across pathogens) generally carry fitness costs when not undergoing treatment. These costs are difficult to measure in vivo, though in vitro fitness-cost estimates are available for select *P. falciparum* genotypes ^31^. Field evidence of fitness costs can appear when resistance decreases are observed after the withdrawal of a first-line therapy, as occurred with chloroquine resistance ^32^. In addition, recent studies on feeding assays between isolated resistant clones suggest that the decrease in relative transmission of artemisinin resistant clones is greater than that modelled here ^33^. The incremental fitness benefits of drug-resistance mutations are equally difficult to parameterize as therapeutic efficacy studies are not powered to measure efficacy on specific genotypes ^34^. A single parameterization of genotype-specific drug efficacies was used in our analysis, even though considerable variation exists for these estimates. A sensitivity analysis on these efficacy values does not alter our general conclusions on partner-drug resistance facilitating artemisinin-resistance emergence (Supplementary Figure 8), but does remind us there is substantial uncertainty in the absolute measures presented in this study, specifically the timelines for future resistance events. Despite this, we were encouraged to find our observed selection coefficients for each ACT fell within the range of selection coefficients estimated in a recent review (Supplementary Figure 5) ^21^. Lastly, model consensus studies are relatively new. Harmonisation and definition of outcome measures needed to be conducted carefully to ensure that the harmonised quantities are not proximally and immediately affecting the outcome measures. Here, we chose the *de novo* mutation rate as our central aligned/harmonized model feature, and we chose a low allele frequency threshold (0.01) to align on to ensure that very few of the detrimental effects of drug resistance would be observed up to this point. As a result the mutation rates chosen and the time to 0.01 may not be representative, however, aligning as we have allowed for comparisons between models for the time till 0.25 frequency. The 0.25 allele-frequency milestone is thus one that is influenced primarily by selection and not mutation, and one that represents an epidemiologically serious and irreversible situation of increasing future drug resistance (unless drug policy is changed).

As a means of basic public health investment, a renewed focus should be placed on early molecular surveillance for partner-drug resistant genotypes. As in all expanded surveillance scenarios, the health-economic equation balancing monetary costs against resistance delays is the crucial one to assess. This general cost-benefit question of investing in drug-resistance surveillance requires a more complete treatment in the malaria literature. Although public health concern typically manifests only after partner-drug resistance is common, the work we present here suggests that early detection of and preemptive action against partner-drug resistance ^35^ would have the benefit of delaying partner-drug resistance, artemisinin resistance, and treatment failure all at once.

## Contributions

OJW, BG, TDN performed the modeling analyses. TN-AT developed genotype-specific efficacy estimation. OJW, BG, TDN, TN-AT, MAP, DLS, LO, RA, and MFB designed the study. All authors evaluated and edited the analysis plan, the results summaries, and manuscript drafts. OJW and MFB wrote the manuscript.

## Acknowledgements

TDN, TN-AT, and MFB are funded by the University of Washington’s Malaria Modeling Consortium grant from the Bill and Melinda Gates Foundation (OPP159934) and by a grant from the Bill and Melinda Gates Foundation (INV-005517). BG and RA acknowledge funding from the Bill and Melinda Gates Foundation (OPP1193472). Thanks to Jennifer Gardy, Erin Stuckey, Scott Miller, Ian Hastings, and Raman Sharma for continuous feedback throughout this collaboration.

## References

1 World Health Organization. Guidelines for the treatment of malaria, 1st edition. 2006 http://www.who.int/iris/handle/10665/149822.

2 Yeung S, Van Damme W, Socheat D, White NJ, Mills A. Access to artemisinin combination therapy for malaria in remote areas of Cambodia. Malar J 2008; 7: 1–14.

3 World Health Organization. WHO briefing on malaria treatment guidelines and artemisinin monotherapies. 2006 https://www.who.int/malaria/publications/atoz/meeting_briefing19april/en/ (accessed Sept 3, 2020).

4 Dondorp AM, Nosten F, Yi P, et al. Artemisinin resistance in Plasmodium falciparum malaria. N Engl J Med 2009; 361: 455–67.

5 Ariey F, Witkowski B, Amaratunga C, et al. A molecular marker of artemisinin-resistant Plasmodium falciparum malaria. Nature 2014; 505: 50–5.

6 Chenet SM, Akinyi Okoth S, Huber CS, et al. Independent Emergence of the Plasmodium falciparum Kelch Propeller Domain Mutant Allele C580Y in Guyana. J Infect Dis 2016; 213: 1472–5.

7 Miotto O, Sekihara M, Tachibana S-I, et al. Emergence of artemisinin-resistant Plasmodium falciparum with kelch13 C580Y mutations on the island of New Guinea. bioRxiv 2019;: 621813.

8 Uwimana A, Legrand E, Stokes BH, et al. Emergence and clonal expansion of in vitro artemisinin-resistant Plasmodium falciparum kelch13 R561H mutant parasites in Rwanda. Nat Med 2020; published online Aug 3. DOI:10.1038/s41591-020-1005-2.

9 Nosten F, White NJ. Artemisinin-Based Combination Treatment of Falciparum Malaria. American Society of Tropical Medicine and Hygiene, 2007.

10 Witkowski B, Duru V, Khim N, et al. A surrogate marker of piperaquine-resistant Plasmodium falciparum malaria: a phenotype–genotype association study. Lancet Infect Dis 2017; 17: 174–83.

11 Nguyen TD, Olliaro P, Dondorp AM, et al. Optimum population-level use of artemisinin combination therapies: A modelling study. The Lancet Global Health 2015; 3: e758–66.

12 Okell LC, Reiter LM, Ebbe LS, et al. Emerging implications of policies on malaria treatment: genetic changes in the Pfmdr-1 gene affecting susceptibility to artemether-lumefantrine and artesunate-amodiaquine in Africa. BMJ Glob Health 2018; 3: e000999.

13 Brady OJ, Slater HC, Pemberton-Ross P, et al. Role of mass drug administration in elimination of Plasmodium falciparum malaria: a consensus modelling study. The Lancet Global Health 2017; 5: e680–7.

14 Penny MA, Verity R, Bever CA, et al. Public health impact and cost-effectiveness of the RTS,S/AS01 malaria vaccine: A systematic comparison of predictions from four mathematical models. Lancet 2016; 387: 367–75.

15 Watson OJ, Okell LC, Hellewell J, et al. Evaluating the performance of malaria genetics for inferring changes in transmission intensity using transmission modelling. Mol Biol Evol 2020; published online Sept 8. DOI:10.1093/molbev/msaa225.

16 Gao B, Saralamba S, Lubell Y, White LJ, Dondorp AM, Aguas R. Determinants of MDA impact and designing MDAs towards malaria elimination. Elife 2020; 9. DOI:10.7554/eLife.51773.

17 Nguyen TD, Tran TN-A, Parker DM, White NJ, Boni MF. Antimalarial mass drug administration in large populations and the evolution of drug resistance. bioRxiv. 2021;: 2021.03.08.434496.

18 Humphreys GS, Merinopoulos I, Ahmed J, et al. Amodiaquine and artemether-lumefantrine select distinct alleles of the Plasmodium falciparum mdr1 gene in Tanzanian children treated for uncomplicated malaria. Antimicrob Agents Chemother 2007; 51: 991–7.

19 Somé AF, Séré YY, Dokomajilar C. Selection of known Plasmodium falciparum resistance-mediating polymorphisms by artemether-lumefantrine and amodiaquine-sulfadoxine-pyrimethamine but not.…. Antimicrob Agents Chemother 2010. https://aac.asm.org/content/54/5/1949.short.

20 World Health Organization. Methods for surveillance of antimalarial drug efficacy. 2009 https://www.who.int/malaria/publications/atoz/9789241597531/en/ (accessed Jan 26, 2021).

21 Hastings IM, Hardy D, Kay K, Sharma R. Incorporating genetic selection into individual-based models of malaria and other infectious diseases. Evol Appl 2020; 13: 2723–39.

22 WorldWide Antimalarial Research Network (WWARN). WWARN Explorer. http://www.wwarn.org/tracking-resistance/wwarn-explorer.

23 Leroy D, Macintyre F, Adoke Y, et al. African isolates show a high proportion of multiple copies of the Plasmodium falciparum plasmepsin-2 gene, a piperaquine resistance marker. Malar J 2019; 18: 126.

24 Inoue J, Silva M, Fofana B, et al. Plasmodium falciparum Plasmepsin 2 Duplications, West Africa. Emerg Infect Dis 2018; 24. DOI:10.3201/eid2408.180370.

25 Bushman M, Morton L, Duah N, et al. Within-host competition and drug resistance in the human malaria parasite Plasmodium falciparum. Proceedings of the Royal Society B: Biological Sciences 2016; 283: 20153038.

26 Hastings IM. Complex dynamics and stability of resistance to antimalarial drugs. Parasitology 2006; 132: 615–24.

27 Ghafari M, Weissman DB. The expected time to cross extended fitness plateaus. Theor Popul Biol 2019; 129: 54–67.

28 Chang H-H, Childs LM, Buckee CO. Variation in infection length and superinfection enhance selection efficiency in the human malaria parasite. Sci Rep 2016; 6: 26370.

29 Hastings IM, Watkins WM. Intensity of malaria transmission and the evolution of drug resistance. Acta Trop 2005; 94: 218–29.

30 Boni MF, Smith DL, Laxminarayan R. Benefits of using multiple first-line therapies against malaria. Proceedings of the National Academy of Sciences 2008; 105: 14216–21.

31 Nair S, Li X, Arya GA, et al. Fitness costs and the rapid spread of kelch13-C580Y substitutions conferring artemisinin resistance. Antimicrob Agents Chemother 2018; 62. DOI:10.1128/AAC.00605-18.

32 Nwakanma DC, Duffy CW, Amambua-Ngwa A, et al. Changes in malaria parasite drug resistance in an endemic population over a 25-year period with resulting genomic evidence of selection. J Infect Dis 2014; 209: 1126–35.

33 Witmer K, Dahalan FA, Delves MJ, et al. Transmission of Artemisinin-Resistant Malaria Parasites to Mosquitoes under Antimalarial Drug Pressure. Antimicrob Agents Chemother 2020; 65. DOI:10.1128/AAC.00898-20.

34 Ljolje D, Dimbu PR, Kelley J, et al. Prevalence of molecular markers of artemisinin and lumefantrine resistance among patients with uncomplicated Plasmodium falciparum malaria in three provinces in Angola, 2015. Malar J 2018; 17: 84.

35 Boni MF, White NJ, Baird JK. The Community As the Patient in Malaria-Endemic Areas: Preempting Drug Resistance with Multiple First-Line Therapies. PLoS Med 2016; 13: 1–7.

