## Supplementary Material for "Pre-existing partner-drug resistance facilitates the emergence and spread of artemisinin resistance: a consensus modelling study"


**Supplementary Material** **1**
Supplementary Methods 2
1 Model Descriptions 2
2 Model Specific Extensions 7
Imperial Model 7
MORU Model 13
PSU Model 18
3 Treatment efficacy on specific genotypes 21
4 Sensitivity Analyses and  Outcome MeasuresScenario Evaluations 23
Supplementary Figures 25
Supplementary Tables 40
Supplementary References 41

### Supplementary Methods

#### 1 Model Descriptions

Three individual-based microsimulations were used to compare patterns of artemisinin-resistance evolution in different contexts of prevalence, treatment coverage, and pre-existing partner-drug resistance. The three models are called (1) the 'Imperial Model', developed at the MRC Centre for Global Infectious Disease Analysis, Faculty of Medicine, Imperial College London; (2) the 'MORU Model', developed at the Mahidol-Oxford Research Unit (MORU), Nuffield Department of Medicine, University of Oxford; and (3) the 'PSU Model', developed at the Center for Infectious Disease Dynamics (CIDD), Department of Biology, Pennsylvania State University. The models all run as daily time-step discrete-event simulations of individuals (humans) who can be infected with *Plasmodium falciparum* malaria and subsequently pass on their infection to other individuals in the simulation. The outputs of model runs and downstream analysis code is publicly available as a reproducible research compendium at https://github.com/OJWatson/art_resistance_consensus_modelling.

An overview of the comparisons between each model are detailed in Table S1. Further methodological details of the individual models and extensions made to previously published models are described below for each model.

**Supplementary Table 1.** Comparative model features

| | **Imperial Model** | **PSU Model** | **MORU Model** |
|---|---|---|---|
| **Key representative publications** | Original Imperial model (Griffin et al. 2016, 2010). Extensions made for modelling parasite genetics in MBE (Oliver J. Watson et al. 2021). | Multiple first line therapies study in Lancet Global Health (Nguyen et al. 2015) and therapeutic efficacy by genotype study bioRxiv preprint (Nguyen et al. 2021)) | Mass drug administration study in eLife (Gao et al. 2020) |
| **Accessibility (Either Github or description of code availability etc.)** | Model code is open source and freely available at www.github.com/OJWatson/magenta . In this study, magenta version 1.3.0 was used (O. J. Watson, Verity, and FitzJohn 2021) | Code is open source at https://github.com/bonilab/malariaibm-MMC-WP2-partnerdrugresistance (version 3.2) | Available on request. |
| **Seasonality** | Yes; not used in present analyses but implemented through changing mosquito abundance. | Yes; not used in present analyses. | Yes; not used in present analyses. |
| **Heterogeneity in exposure** | Yes | Yes | Yes |
| **Blood-stage parasite densities modelled** | Not explicitly. Parasite densities are approximated by tracking a parasite in terms of the infection status that the parasite causes the human in the absence of superinfection (symptomatic, asymptomatic, sub-patent). These states are then used to calculate the relative probability of different strains being passed on when a human infects a mosquito. See (Oliver J. Watson et al. 2021) for further detail. | Yes | No |
| **Parameterization for clinical incidence** | Fitted to cross-sectional age-incidence data from 23 sites in Africa capturing differences between active and passive case detection (Griffin, Ferguson, and Ghani 2014). | Calibrated to data sets assembled for five different African studies measuring age-specific clinical incidence and EIR. No formal model fitting done. | We parameterised (1) the age dependent risk of infection during the first 10 of life was estimated from data from 8 endemic countries in sub-Saharan Africa (Aguas et al., 2008). Given (1), clinical incidence was calibrated to a dataset of immunity markers collected in 4 Cambodian sites, that describes seroconversion rates over time and extrapolates age profiles of clinical immunity (Cook et al. 2012) |

| | | | |
|---|---|---|---|
| **Parameterization for severe disease and mortality incidence** | Severe disease model (Griffin et al. 2015) fitted to data from northern Tanzania (Reyburn et al. 2005) and to severe disease vs. prevalence relationship from data of multiple sites (Marsh and Snow 1999). Mortality due to malaria is based on Africa-wide data from verbal autopsy and parasite prevalence (Rowe et al. 2006). | Mortality only. No tracking of severe malaria. | Subdivides population into non-immune & immune classes and clinical and subclinical infections. Probability of clinical outcome is defined as: <br><br> Probclin = (imm_a*exp(−0.1(cml−1)) + exp(−imm_b*(cml+1)))/sqrt(lvl) <br><br> Where (*cml*) is the cumulative number of infections and (*lvl*) is the mean immunity level for each age category. The parameter set θ={imm_a,imm_b} was estimated from Cambodian data (Cook et al. 2012) |
| **Drug interventions – PK- PD Drug action etc** | Slow clearance, treatment failure incorporated. PKPD not explicitly incorporated in transmission model. PKPD are modelled separately and their outputs incorporated as parameters that determine the probability of slow parasite clearance or late parasitological failure as defined in (Slater et al. 2016). | Single compartment PK model used, with daily parasite killing, as a function of drug concentrations and parasite genotypes, for PD model. | Includes slow clearance, treatment failure, appearance and spread of resistance. Parasite clearance modelled using PD data from parasite clearance studies, with daily parasite killing, as a function of initial drug concentration, drug IC50 and parasite genotype. |
| **Genotype tracking and sensitivity to drugs** | Yes, four key resistance loci and two copy number variants tracked. 64 total genotypes. | Yes, four key resistance loci and two copy number variants tracked. 64 total genotypes. | Yes, four key resistance loci and two copy number variants tracked. 64 total genotypes. |
| **Vector control Interventions** | LLIN, IRS, Larval control (larviciding & pupaciding), and ivermectin. | Indirect vector control only, via transmission parameter that determines the daily amount of biting. | Yes – LLIN, IRS, Larval Control |
| **Treatment interventions** | Treatment of clinical disease and severe disease, by specified drug and diagnostic. <br> Mass screen and treat, IPTi, IPTc/SMC and IPTp/IST for separate pregnancy model | Yes, many types of drug policies, private-market drug sales, and mass drug administration. | Yes treatment of clinical disease, MDA, MSAT, adjuctive primaquine, TME, iPT |
| **Treatment seeking and drug coverage** | Explored in the sensitivity analysis | Explored in the sensitivity analysis | Explored in the sensitivity analysis |
| **Spatial dynamic model** | Present analysis run in a single location. | Present analysis run in a single location. | Yes, but run at admin level 1 for these analyses. |
| **Super-infections, co-infections, multiplicity of infection** | Multi-strain models track multiplicity of infection. Individuals can be infected by a single mosquito with multiple sporozoites with an assumed geometric mean | Yes, tracked explicitly as coinfections arising only from superinfection events. | Yes, each human individual can carry up to 10 different parasite populations (one per inoculum). Each inoculum is considered to be a clonal population upon emergence |

| | | | |
|---|---|---|---|
| | of 10 parasites passed on during an infectious bite, with 21% surviving to produce a liver stage infection (21% parameterised in (Oliver J. Watson et al. 2021)), which means multi-clonal infections can arise from a single bite. Different strains | | from the liver, with one parasite acquired from an infectious bite. |
| **Heterogeneity in Exposure** | Exposure varies both by age and between individuals. | Exposure varies both by age (v3.0.4) and between individuals. | Exposure varies both by age and between individuals. |
| **Duration of infection** | Duration of infection is "Erlang-like" distribution (convolution of exponential distributions), with overall duration calibrated to malariatherapy data. | Not drawn from a predetermined distribution. The duration of infection is determined by explicit modeling of parasitaemia and how drugs and the immune system act on parasites. Calibrated to malariatherapy data. Durations of infection in the model range from 60 to 281 days. | Asymptomatic infection duration based on malariatherapy data and clinical follow-up data from endemic areas, as well as drug efficacy data (PD). Also estimated from a set of 8 Endemic areas from Sub-Saharan Africa (Aguas et al 2008). |
| **Clinical disease and history of exposure** | A proportion of infected individuals go on to develop clinical disease (parameterised in (Griffin, Ferguson, and Ghani 2014)). Immunity to clinical disease develops with exposure and age, and also has a maternally acquired component. | A proportion of infected individuals go on to develop clinical disease. Immunity to clinical disease develops with exposure and age, and also has a maternally acquired component. | Proportion developing clinical disease depending on cumulative immunity from prior exposures and immunity level (indicator of recent exposure). |
| **Decay of natural immunity** | Exponential decay of naturally acquired immunity (parameterised in (Griffin, Ferguson, and Ghani 2014)). | Exponential decay of naturally acquired immunity. | Exponential decay of naturally acquired immunity as estimated in (Aguas et al. 2008) |
| **Infectiousness and gametocyte models** | Human infectiousness to mosquitos is dependent on the infection status of the human (clinical, asymptomatic, sub-patent) and if asymptomatic also determined by their immunity determining their probability of being detected by microscopy (parameterised in (Griffin, Ferguson, and Ghani 2014)). | Human infectiousness to mosquitos is a function of asexual parasite density, with a time lag built in to mode the fact that infectious gametocytemia lags asexual parasitaemia.. | Infectiousness depends on lagged development of sexual stage parasites and seasonal transmission equation from fitting to incidence data. It is informed by parasite density in an indirect way: clinical individuals are assumed to have a higher mean infectiousness compared to asymptomatic individuals as they carry higher parasite density loads. |
| **Entomological models** | Vector control interventions modelled through altering the life expectancy of a mosquito, altering the rate of anthrophagy. (described and parameterised in (Griffin et al. 2016)). | 11-day lag built in so that FOI today depends on the biting done by mosquitoes 11 days ago. No other entomological features built in. | Full IBM component to mosquito dynamics. For this exercise we use a simplified version where only infectious mosquitoes are tracked individually. . |
| **Recombination model** | Multiple parasites taken up by mosquito to form *n* | Recombination can occur in a mosquito bite on a multi-clonal | No recombination. No interrupted feeding. |

| | | | |
|---|---|---|---|
| | oocysts, which undergo recombination to yield up to 4xn genotypes of sporozoites in the mosquito. No interrupted feeding by single mosquito on multiple humans. | host. Parasites are taken up by the mosquito in proportion to their parasite density. A full recombination table is built for all possible forces of infection resulting from this host's contribution to the next generation of infectious sporozoites, according to the normal rules of Mendelian genetics. No interrupted feeding allowed in present analyses. | |
| **Mutation model** | Back mutation present. Mutation can occur during treatment and in the absence of treatment. 1 rate for both calibrated during the calibration exercise. | No back mutation. Mutation can occur during treatment only when the mutation confers a resistance benefit to the current treatment. | No back mutation. Mutation can occur during treatment (higher rate), and in the absence of treatment (lower rate). The higher rate was calibrated during the calibration exercise. |

#### 2 Model Specific Extensions

**Imperial Model**

The base model has been extensively described in (Oliver J. Watson et al. 2021). Briefly, it is an individual based, discrete time, stochastic model, with individual mosquitoes explicitly modelled, originally developed for characterising the impact of transmission intensity on neutral genetic diversity. Below, we detail the extensions made for simulating antimalarial resistance.

##### Parasite genetic barcode extensions for modelling resistance

In order to model antimalarial resistance, we have adapted the parasite barcode used in Watson et al. for the simulation of resistance (Oliver J. Watson et al. 2021). In the new formulation, each position in the barcode represents either the absence (barcode position is equal to 0) or presence (barcode position is equal to 1) of a resistance mutation associated with resistance to a particular drug. For example, Supplementary Table 2 shows how a barcode with three loci can be used to represent resistance to DHA-PPQ and ASMQ.

**Supplementary Table 2. Example barcode alterations to model antimalarial resistance**

| Artemisinin Resistance | Piperaquine Resistance | Mefloquine Resistance | Phenotype |
|---|---|---|---|
| 0 | 0 | 0 | Wild type parasite. Fully susceptible. |
| 1 | 0 | 0 | Resistant to artemisinin |
| 0 | 1 | 0 | Resistant to PPQ |
| 0 | 0 | 1 | Resistant to MQ |
| 1 | 1 | 0 | Resistant to DHA-PPQ |
| 1 | 0 | 1 | Resistant to ASMQ |
| 0 | 1 | 1 | Resistant to PPQ and MQ |
| 1 | 1 | 1 | Multidrug resistant to DHA-PPQ and ASMQ |

Barcodes are used to track populations of parasites that are introduced from an infectious mosquito bite. We assume loci are genetically unlinked ( genes known to confer resistance to the five first-line ACTs recommended by the WHO each occurring on different chromosomes) and consequently segregate independently during recombination.

**Fitness Costs**

Antimalarial resistance is assumed to introduce a fitness cost to resistant parasites compared to wild type parasites. Fitness costs are associated with each resistance locus and are assumed to be multiplicative. The fitness cost manifests as a reduction in parasite density that is assumed to reduce the probability of the resistant parasite being passed on to a mosquito. This can be expressed mathematically as follows. Let $b = \left[b_1, b_2, ..., b_m\right]$ describe the vector of barcode loci for a barcode of length $m$. For example, the ASMQ resistant strain in Supplementary Table 2 is represented by the vector $[1,\ 0,\ 1]$. If $\mathrm{v}_j$ is the resistance cost associated with barcode locus $j$, then the comparative fitness cost due to resistance, $r$, for the given parasite is simply:

$$r = 1 - \prod_{i=j}^{m} \mathrm{v}_j$$

In this study, the wild type allele at each locus Is assumed to have no fitness cost ($v_j = 1$) and thus $r = 1$ for the wild type parasite in Supplementary Table 2. In this analysis, each resistance locus is assumed to have a fitness cost equal to 0.0005, i.e. $v_{i...m} = 0.0005$. As a result, the relative fitness of a genotype with $k$ resistance mutations, in the absence of any drugs present in the blood, is $(1\ -\ 0.005)^k$.

The number of oocysts generated from each mosquito bite on an infectious human is drawn from a zero-truncated negative binomial distribution with mean = 2.5 and shape = 1. The selection of which parasite strains from the infected individual contributed to the oocysts is given by the relative probability that a given strain will be chosen in an individual with $n$ gametocytogenic parasite strains, and is given by:

$$cr = \left[c_1 r_1,\ c_2 r_2, ...c_n r_n\right]$$

$r_i$ is the fitness cost associated with parasite strain $i$ and $c_i$ is the contribution of parasite $i$ to onward infection, which will be either $c_T$, $c_D$, $c_A$ or $c_U$ depending on the infection status of strain $i$, denoted here as $X_i$. The probability that an infected individual infects a mosquito is still determined by the set of parameters determining the onward contribution to transmission, $\{c_T, c_D, c_A, c_U\}$, which are based on the infection status of the individual, denoted here as $Y$. However, if the strains responsible for the human's current infection state, i.e. all strains that match the human's infection state, are resistant then the probability of onward transmission is determined by the highest onward contribution of these strains, which is given by:

$$max\{cr_i : X_i = Y\}$$

In this way, fitness costs both affect the relative probability that a resistant strain is transmitted compared to a wild type strain in a mixed infection, while also reducing the probability that transmission occurs in individuals where the highest parasite density strain is resistant.

**Clinical and Treatment Outcomes**

With the addition of resistance, the treatment efficacy now varies and is determined both by the genotype of the parasite strains, the parasite density of each strain and the drug used to treat the infection. The probability that any given strain is cleared by drug $z$ can be expressed as $ez_{\beta(b)}$, where $\beta(b)$ is an adapted conversion of binary to decimal integers and is given by:

$$\beta(b) = \left(\sum_{j=1}^{m} b_j . 2^{j-1}\right) - 1$$

Using the same representation of a parasite genotype as in Supplementary Table 2 the efficacy of drug $z$ can be expressed by the vector $ez = \left[ez_1, ez_2, ..., ez_{2^m}\right]$ for simulations in which the number of loci being modelled is equal to $m$. $ez_1$ represents the efficacy of the drug against the wild type parasite, i.e. the barcode vector $b$ represented by a vector of length $m$ filled with zeros. Importantly, $ez$ reflects the probability that the drug will clear a parasite that has led to a symptomatic infection, i.e. the parasite strain is at a sufficiently high parasite density to trigger symptoms and seek treatment. This is equivalent to the probability of successfully clearing a symptomatic infection such that the infection does not recrudesce and lead to a 28-day treatment failure.

In this analysis, we used the drug efficacy by genotype table parameterised in (Nguyen et al.

2021), which was used by each group to parameterise the efficacy of each drug on each resistance genotype. 64 genotypes are included by allowing for variation at the K76T locus in *pfcrt*, the N86Y and Y184F loci in *pfmdr1*, the C580Y locus in pfkelch13, copy-number variation (CNV) of *pfmdr1*, and CNV of the *plasmepsin-2,3* genes. CNV is only separated into 'single copy' or 'multiple copies'. See Section 3 - Treatment efficacy on specific genotypes for more information about this. In the Imperial model, the efficacy of each drug determines the probability of a 28-day parasitological failure, with the probability of late parasitological failure for each parasite genotype modelled shown in Supplementary Table 3.

Parasites from previous infections are assumed to be at a lower parasite density than the infecting strain that triggered the clinical infection (and hence why the individual is seeking treatment) and will be more likely to be cleared by the drug. In the model, the infection state of each strain, $X_i$, as well as the day the strain was acquired, $t_0$, and the time the strain will move out of the infection state, $t_1$, is tracked. This information is used to define the probability that strain $i$ will recrudesce after treatment, $P(Recrudesce)_i$, which is given by:

$$P(Recrudesce)_i = \begin{cases} ez_{\beta(b_i)}\,, & X_i = Y \\ ez_{\beta(b_i)}\left(\dfrac{t_1 - t_c}{t_1 - t_0}\right), & X_i = A \\ 0\,, & X_i = U \end{cases}$$

where $t_c$ is the current time. $P(Recrudesce)_i$ assumes that parasites below ~200p/μl (state U) will always be cleared regardless of the parasite phenotype. $P(Recrudesce)_i$ also assumes that the probability that an asymptomatic parasite above ~200p/μl (state A) will recrudesce is linearly related to the age of the infection and is at its highest when it first enters state A.

The probability that a treated infection will be successfully cleared by drug $z$, $P(Cleared)$, is equal to one minus the highest probability of a strain recrudescing, which is given by:

$$P(Cleared) = 1 - max\{P(Recrudesce)_i : 0 \le i \le n\}$$

This assumes that the multiplicity of infection does not directly affect the probability that an individual will be cleared, i.e. only one Bernoulli trial with probability $P(Cleared)$ is used to determine if all parasite strains were cleared.

If an individual fails treatment, it is assumed that they will recrudesce to yield a late parasitological failure (LPF) and move into state A after the prophylactic period of the drug has finished. Whether each parasite strain in a multiply infected individual recrudesces during a LPF is dependent on $P(Recrudesce)_i$. Bernoulli trials are conducted for each parasite strain except for one random strain for which $P(Recrudesce)_i = P(Cleared)$, which ensures that one of the most likely strains to recrudesce did actually recrudesce and cause a LPF.

If an individual successfully clears all parasites, they will either move directly into a state P or they will remain in the treated compartment for a longer duration resulting from slow parasite clearance (SPC). SPC is assumed to always occur if any of the pre-treatment strains that contributed to the clinical disease were resistant to any component of the drug. The duration of SPC was set equal to 10 days based on previous modelling studies estimating parasite clearance rates associated with SPC (Slater et al. 2016). During SPC it is assumed that all parasite strains not resistant to the drug given have cleared and will thus not contribute to onward infection during SPC.

Lastly, individuals in state P can be reinfected before returning to state S, depending on the genotype of the infecting strain, how recently they were treated and the ACT used, reflecting the different half-lives of available partner drugs. For all drugs the probability of reinfection in state P increases as the partner drug wanes. As described in (Bretscher et al. 2019), we use the gamma distribution, $\Gamma(\alpha_i, \beta_i)$, where $\alpha_i$ and $\beta_i$ are the shape and rate parameter respectively of the Gamma distribution, for drug $i$, to describe the probability of reinfection in individuals treated with AL and ASAQ. We use $\alpha_{AL} = 93.5$ and $\alpha_{ASAQ} = 16.8$ for the shape parameters for AL and ASAQ. These values represent the mean posterior estimate for the shape parameter in (Bretscher et al. 2019), which were estimates across a number of trial sites in Africa. We use $\beta_{AL} = 5.22$ and $\beta_{ASAQ} = 0.94$ for the rate parameters, which yield mean durations of protection equal to 17.9 days and 17.8 days respectively. These durations are longer than the posterior mean duration of protection estimated in (Bretscher et al. 2019) and more closely reflect study sites in (Bretscher et al. 2019) with the longest times to reinfection for AL and ASAQ. This was chosen because the mean duration was estimated across study sites with known resistance markers and subsequently we chose to match to relationships in sites that were assumed to have the least amount of partner drug resistance for each drug. For DHA-PPQ, we assume the probability of reinfection is described by a Weibull survival curve, with scale and shape equal to 28.1 and 4.4 respectively, as estimated in (Okell et al. 2014) , with a mean duration of protection equal to 25 days. The described periods of prophylaxis are used to define the per-day probability of an individual being reinfected if the parasites introduced during an infectious bite are not resistant to the partner drug. However, if any of the introduced parasites are resistant to the partner drug, we assume a shorter period of prophylaxis. For AL and ASAQ, we use $\beta_{AL} = 10.75$ and $\beta_{ASAQ} = 1.45$, resulting in mean durations of protection equal to 8.7 days and 11.6 days respectively. These shorter durations were chosen to match to prophylactic profiles in sites in (Bretscher et al. 2019) with the shortest durations of protection due to the presence of partner drug resistance. For DHA-PPQ, we used a larger scale parameter of 59 for the Weibull survival curve, approximately halving the mean duration of protection to 12.4 days.

In order to simulate individual level variation in drug clearance and prophylaxis we first fit a Hill function to relate the assumed exponential clearance of the partner drug to the described period of prophylaxis for each drug on both wild type and resistant parasites. For each drug, we assume the mean drug lifetime is equal to $ln(2)$ * mean duration of prophylaxis of each drug against wild-type parasites (17.9, 17.8 and 25 days for AL, ASAQ and DHA-PPQ respectively from earlier prophylaxis fitting). When an individual moves from being treated to being in a state of prophylaxis, we draw the time at which they will become fully susceptible again from an exponential distribution with mean equal to the mean duration of prophylaxis for each drug. During this prophylactic period, we assume the individual’s drug concentration decays exponentially such that they have fully eliminated the partner drug when returning to state S, with their drug concentration on each day used to calculate the probability of early reinfection using the earlier fitted Hill functions.

#### MORU Model

The base model has been extensively described in (Gao et al. 2020). Briefly, it is an individual based, discrete time, spatially explicit, stochastic model, with mosquito population dynamics as well as human population movement. The spatial features have been turned off for the purposes of the analyses presented here.

##### Population

As mentioned in the original model publication, since the vast majority of bites (at least for the vector species considered here) occur overnight, we are solely concerned with night time mosquito biting patterns and human behavior. In general we would consider that daily population flows between villages are best characterised by a modified gravity model, but given that the models presented here were calibrated for different geographical settings and have different spatial resolutions and inherent human and mosquito mobility formulations, we chose to remove the human movement component of our model. Thus, to ensure that potential discrepancies between model predictions presented here are not driven by the significant differences in the mobility and spatial resolution components of the models, we assumed a closed population of 100,000 individuals, which in our framework is equivalent to discriminating a single village containing all humans and mosquitoes as described below.

##### Transmission

Transmission within a village follows a quasi-homogeneous process whereby each mosquito is equally likely to bite any individual. Given that people can move freely within their village whilst mosquitoes are actively seeking and biting, we can reasonably assume that all humans in a given village can potentially be bitten by any one mosquito in that village.

Malaria transmission is spatially heterogeneous (Erhart et al. 2005; Cui et al. 2012; Gryseels et al. 2015) as manifested by significantly different disease incidence rates within countries, but entomological data to inform the distribution of mosquito densities at scale is lacking. Due to this lack of mosquito abundance data and the fact that each model was calibrated for different settings where mosquito bionomics are likely to vary, and thus to minimise potential divergence across models, we assumed that humans and mosquitoes are enclosed in a single village of 100,000 individuals.

**Immunity and symptomatology**

Each human agent is assigned two properties in relation to immunity, namely Cumulative Number of Exposures and Immunity Level. Both properties are set to 0 for newborns. The likelihood of clinical symptoms brought on by a single infection is given by:

$$clinical_{prob_n} = e(-\ 0.15(moi_n - 1))\frac{0.1e(-0.1cml_n - 1)\ + e(-0.9cml_n)}{lvl_n^{\ 0.5}}$$

where *moi*, *cml* and *lvl* denote the multiplicity of infection, cumulative exposure to malaria, and immunity level properties of the human agent, respectively. Although we only increase the immunity level if individuals resolve their infection (presumably due to increased antibody killing activity), the cumulative exposure is updated with each infectious bite received. Thus, individuals can accrue some immunity with superinfections. One level of clinical immunity is gained by a human agent every time his infection list is emptied. Immunity loss starts 40 days after one level of immunity is gained. Immunity is lost at a rate of 60 days$^{-1}$. Therefore, each human agent at immune level one is clinically immune an average of 100 days. A loss in immunity prompts a reduction in immunity level and not the immune status per se.

Non-clinical (asymptomatic) untreated infections are assumed to last an average of 160 days, following data from the malaria therapy experiments (Collins and Jeffery 1999).

**Interventions**

The base model includes a range of possible malaria control interventions, from Insecticide Treated Nets (ITNs/LLINs) to more extreme and logistically complex ones like Targeted Malaria Elimination (TME). For this model comparison exercise, we decided to remove all control interventions except baseline clinical malaria management. This is the foundation of any malaria control program as it can be parameterised to reflect the percentage of clinical infections that receive appropriate treatment (antimalarial drugs). The choice of drugs dispensed is made explicit in each of the presented results.

**Genotype to phenotype map**

In our base model, each infection is treated as an individual entity with its own life cycle within the blood system of a human agent, and its drug-resistance profile is a product of the genetic information in the six alleles conferring resistance to the drugs considered here. Genotype-specific drug efficacies are described in Section 3 - Treatment efficacy on specific genotypes.

**Within host dynamics**

***PK/PD***

We use a simple 1-compartment pharmacokinetic (PK) model that assumes drug concentration drops below a pre-defined minimum inhibitory concentration (MIC) following an exponentially distributed rate. The MIC defines the point in time after treatment at which the drugs administered are no longer able to counteract parasite growth, i.e. the day after which the infection can only be cleared by immunity. Our PK compartment does not reflect drug concentration, but rather a drug action status defining the potential parasite clearance by drug action. If the drug action status is ON, i.e. drug concentration is above the MIC, drug killing effects for each drug considered are implemented as a daily probability of clearance that results in the expected treatment failure rate at day 28 for each combination of drug given and parasite genotype. We assumed the mean times to reach MIC are 25 days for amodiaquine, 14 days for lumefantrine, and 30 days for piperaquine. Artemisinin-derivatives are assumed to only be present at the day of dosing.

Parasite killing rates depend on the person's transmission status (*s*), with parasite clearance in not yet infectious people generally slower than that in individuals carrying gametocytes. Clearance of parasites with drug resistance phenotype *h* by drug *d* then follows:

$$Clearance_{shd} \sim B(N_{shd}, c_{shd})$$

where $c_{shd}$ is an element of a 3-dimensional matrix *C* of size |S|*|H|*|D| and *B* denotes a binomial distribution. $N_{shd}$ determines the number of parasite sub-populations affected by $c_{shd}$, thus a human host is only declared cleared from infection once every parasite sub-population has been cleared from their bloodstream. The parameter values in $c_{shd}$ have been calibrated to produce the expected treatment failure rates in Section 3, as per the common genotype-specific drug efficacies table, given the drug PK described above.

***Mutation***

Per parasite subpopulation, we consider a daily probability of mutation at any of the alleles conferring resistance to either artemisinin or the partner drugs. These mutations are considered independent events meaning they may happen on separate days or on the same day, and there is no precedence between the two events. A base mutation rate of $10^{-9}$ is used in the absence of drug pressure. That rate is five orders of magnitude higher when drugs are present in the blood system (Pongtavornpinyo et al. 2009).

***Clone selection***

Our base model allows for co-infection with a large number of clones (defined as parasite populations originating from different inocula, regardless of genotype). In the version used here, we simplify within-host dynamics by imposing a set of rules to determine the likelihood of fixation of different parasite clones under different drug pressure contexts. Given the assumed differences in parasite fitness according to genotype, we determine that: *i*) in the presence of combination therapy, double resistant parasites are likely to fix if present, followed by single resistant genotypes. Thus, if the parasite population is a mix of single mutants and wild-type parasites, the single mutant parasites will eventually outcompete the wild-type parasites under drug pressure, will form the bulk of mature gametocytes, and subsequently be transmitted to mosquitoes upon a bite. If there is a mix of double mutants with any other genotype subpopulation, the double mutant population will fix. *ii*) when there are no drugs in the bloodstream, a random draw decides which parasite subpopulation will be transmitted to the mosquito. This effect tries to account for infections where the drug-resistance allele emerges at different infection time points, infections where drugs were given at subtherapeutic levels, and infections with a greater multiplicity of infection, and is re-evaluated over the course of the infection, upon a new mosquito bite. Importantly, transmissibility of different clones to mosquitoes is obviously dependent on relative parasite frequencies of different clones, which are a direct result of the dynamic processes described above. Using an expanded version of an in-house within-host *P. falciparum* model (Saralamba et al. 2011), we determined that there is a critical time window for the fixation of resistant parasite populations (Supplementary Figure 13) Independently of when the resistance clones emerge (middle panel), fixation seems to always occur at day 5-7 (right-side panel) post drug treatment. This finding allowed us to set a six-day lag for the higher mutation rate to be applied. This means that for the first six days after treatment is initiated, the base mutation rate is used; if after six days there is still drug present in the blood, the higher mutation rate is applied. Once there is no drug left in the blood, the mutation rate reverts to the base rate if the infection has not yet been cleared. We thus are able to reproduce the complex within host dynamics generated by far more complex models by modulating the use of the mutation rates and asserting fixed allelic dominance functions that determine which parasite clones will fix under different drug pressure scenarios. We do not model back mutation events.

#### PSU Model

Model characteristics are described in detail in (Nguyen et al. 2015, 2021), and summarized below with changes noted.

##### Population

A population of 100,000 individuals is modeled. Individuals can be uninfected, or infected with one or more clones of *Plasmodium falciparum*; each clone is its own genotype. In low-transmission regions, the vast majority of infections are single-clonal, while in high-transmission regions mult-clonal infections are common. The model's distribution of clones per person is calibrated to data in (Owusu-Agyei et al. 2002) from a high-transmission region of northern Ghana. Mosquito bites occur on individual hosts in the model determined by each person's 'biting attractiveness' parameter. Biting attractiveness is assigned at birth in the model, and is drawn from a gamma distribution with a coefficient of variation of 2.0 (Smith et al. 2005). An extrinsic incubation period of 11 days is modeled, and new infections are generated every day in the model based on genotypes that were sampled by mosquitoes 11 days ago. Genotypes distributions drawn 11 days ago are adjusted to account for recombination occurring in hosts with multi-clonal infections, with a standard recombination table. In other words, a 64 x 64 x 64 table is maintained showing the probability that genotype A recombining with genotype B would form genotype C. The within-host frequencies of genotypes A and B give the probability of sampling both genotype A and B, and the value in the table gives the probability that the offspring is genotype C. A final genotype distribution -- based on the frequencies of currently circulating genotypes, and the probabilities that their offspring after recombination are "genotype C" -- is used to determine onward infection to new hosts.

##### Parasitaemia and Immunity

Each clone in each host has its own blood-stage parasite density variable which influences transmission probability to mosquitoes, according to (Ross, Killeen, and Smith 2006). Gametocyte density is assumed to be proportional to asexual density, and gametocyte dynamics are not modeled separately; however a 'delayed version' of the asexual-density variable is used to determine onward transmission as rises and falls in gametocyte densities mostly follow those in asexual densities. Parasitaemia in hosts goes up after an infectious bite that results in symptoms, with peak parasite density at the point of fever/presentation drawn randomly to be between 2000 parasite/μl and 200,000 parasite/μl (uniform distribution on $\log_{10}$-scale, between 3.3 and 5.3). An infectious bite that does not result in symptoms will start with a mean parasitaemia given by $10^x$, where x is drawn from a normal distribution with mean 3 and standard deviation 0.5. This gives a mean starting parasitaemia level of 1000/μl with 95% of draws giving a parasite density between 100/μl and 10,000/μl for asymptomatic individuals.

**Duration of Infection**

Asymptomatic infections are cleared according to a host's immune level *M*, which is simply a relative indicator between 0.0 and 1.0 of minimum and maximum clearance rates by the immune system. Daily clearance rates by the immune system occur at the rate:

$$D_{t+1} = w_R \cdot [0.8036(1 - M_t) + 0.9572M_t] \cdot D_t$$

and are calibrated (as explained in supplement section 1 of (Nguyen et al. 2015)) to several other well-known data sets and approaches ((Filipe et al. 2007; Molineaux and Gramiccia 1980; Maire et al. 2006; Eyles and Young 1951)). Infections are a minimum of 60 days long and a maximum of 281 days long.

**Symptomatic Infection**

Upon an infectious bite, a host will progress to a case of symptomatic febrile malaria with probability

$$P_{clin} = \frac{0.99}{1 + (M / 0.4)^4}$$

and with probability 1 - $P_{\text{clin}}$ the parasites from the new bite will expand to a parasite density between 100/μl and 10,000/μl and establish as an asymptomatic infection. M in the above equation refers to the host's current level of immunity. This calibration was done to several clinical-incidence-by-age data sets from Africa (section 11 of supplement, (Nguyen et al. 2015))

**PK/PD**

A 1-compartment pharmacokinetic (PK) model is used with basic exponential decay modeling drug clearance. Half-lives used are 9 days for amodiaquine, 4.5 days for lumefantrine, and 28 days for piperaquine. Half-lives for artemisinin derivatives are not modeled as they are very short-lived and the simulation has a one-day time-step. Artemisinin-derivatives are simply “present” in the blood on the day of dosing, absent the day after dosing (if no other dose is taken), and are modeled to have a daily $p_{max}$ = 0.999 on drug-sensitive parasites, which translates to a daily killing rate of 0.9982 for an average drug concentration.

**Fitness costs**

All single-resistant parasites are assumed to have a daily fitness cost $c_1$ = 0.0005, which translates to an annual fitness cost of 0.167, i.e. a relative fitness of 0.833. Each resistance mutation (i.e. non-wild-type) adds an additional and equal fitness cost, and fitness costs are multiplicative and independent. In other words, the relative fitness of a genotype with $k$ resistance mutations, in the absence of any drugs present in the blood, is $(1 - c_1)^k$.

#### 3 Treatment efficacy on specific genotypes

All three models (PSU, MORU, Imperial) use the same locus-based drug-resistance model and genotype-specific drug efficacies. 64 genotypes are included by allowing for variation at the K76T locus in *pfcrt*, the N86Y and Y184F loci in *pfmdr1*, the C580Y locus in pfkelch13, copy-number variation (CNV) of *pfmdr1*, and CNV of the *plasmepsin-2,3* genes. CNV is only separated into 'single copy' or 'multiple copies'. With 4 drugs used in the simulation -- artemisinin derivatives, lumefantrine, amodiaquine, piperaquine -- a total 64 x 4 = 256 drug-genotype interactions need to be modeled as an individual can be infected with any genotype and treated with any combination therapy. The monotherapy parameters need to be input into the simulation as well in case an individual is bitten and infected during a time window when only one residual drug is present in the blood. Each of the 256 drug-genotype combinations is parameterized by a $p_{max}$ value (maximum killing rate) and an EC50 value (concentration at which killing power is reduced by half). These values are calibrated in a single-compartment PK/PD model to obtain efficacies that match the clinical trial literature, with some inference and imputation on what genotypes would have been circulating at the time each clinical trial was run. In the table below, a total of 64 x 3 = 192 efficacies are presented for three ACTs and 64 genotypes. See Supplementary Appendix 2 in (Nguyen et al. 2021) for full details as well as 28-day on calculation of treatment efficacies of each ACT modelled against each genotype in Supplementary Table 3.

In Supplementary Table 3 (below), the 6-character genotype code (e.g. KNY1C1) is, from left to right: K76T, N86Y, Y184F, CNV of pfmdr1, C580Y, CNV of plasmepsin-2,3 genes. To highlight the contribution of specific alleles towards treatment efficacies, we constructed a random forest model to predict treatment efficacy based on the presence/absence of specific alleles (Breiman 2001). The importance of each allele in the random forest based on the mean decrease in the Gini index is shown in Supplementary Figure 14 with the predictions of the random forest.

#### Supplementary Table 3: ACT 28-day probability of parasite clearance

| Genotype | DHAPPQ | ASAQ | AL |
|---|---|---|---|
| KNY1C1 | 0.972486 | 0.962073 | 0.915158 |
| TNY1C1 | 0.972486 | 0.947158 | 0.929358 |
| KYY1C1 | 0.972486 | 0.905281 | 0.953453 |
| TYY1C1 | 0.972486 | 0.891124 | 0.964582 |
| KNF1C1 | 0.972486 | 0.976982 | 0.889945 |
| TNF1C1 | 0.972486 | 0.958879 | 0.908054 |
| KYF1C1 | 0.972486 | 0.930465 | 0.915158 |
| TYF1C1 | 0.972486 | 0.917383 | 0.929358 |
| KNY2C1 | 0.972486 | 0.962073 | 0.858645 |
| TNY2C1 | 0.972486 | 0.947158 | 0.869483 |
| KYY2C1 | 0.972486 | 0.905281 | 0.896817 |
| TYY2C1 | 0.972486 | 0.891124 | 0.915158 |
| KNF2C1 | 0.972486 | 0.976982 | 0.830015 |
| TNF2C1 | 0.972486 | 0.958879 | 0.844322 |
| KYF2C1 | 0.972486 | 0.930465 | 0.858645 |
| TYF2C1 | 0.972486 | 0.917383 | 0.869483 |
| KNY1Y1 | 0.928664 | 0.895737 | 0.795336 |
| TNY1Y1 | 0.928664 | 0.864359 | 0.828863 |
| KYY1Y1 | 0.928664 | 0.771709 | 0.877727 |
| TYY1Y1 | 0.928664 | 0.735268 | 0.907645 |
| KNF1Y1 | 0.928664 | 0.948359 | 0.723351 |
| TNF1Y1 | 0.928664 | 0.892625 | 0.753201 |
| KYF1Y1 | 0.928664 | 0.824725 | 0.795336 |
| TYF1Y1 | 0.928664 | 0.794297 | 0.828863 |
| KNY2Y1 | 0.928664 | 0.895737 | 0.646288 |
| TNY2Y1 | 0.928664 | 0.864359 | 0.684442 |
| KYY2Y1 | 0.928664 | 0.771709 | 0.75155 |
| TYY2Y1 | 0.928664 | 0.735268 | 0.795336 |
| KNF2Y1 | 0.928664 | 0.948359 | 0.570356 |
| TNF2Y1 | 0.928664 | 0.892625 | 0.604481 |
| KYF2Y1 | 0.928664 | 0.824725 | 0.646288 |
| TYF2Y1 | 0.928664 | 0.794297 | 0.684442 |
| KNY1C2 | 0.768484 | 0.962073 | 0.915158 |
| TNY1C2 | 0.768484 | 0.947158 | 0.929358 |
| KYY1C2 | 0.768484 | 0.905281 | 0.953453 |
| TYY1C2 | 0.768484 | 0.891124 | 0.964582 |
| KNF1C2 | 0.768484 | 0.976982 | 0.889945 |
| TNF1C2 | 0.768484 | 0.958879 | 0.908054 |
| KYF1C2 | 0.768484 | 0.930465 | 0.915158 |
| TYF1C2 | 0.768484 | 0.917383 | 0.929358 |
| KNY2C2 | 0.768484 | 0.962073 | 0.858645 |
| TNY2C2 | 0.768484 | 0.947158 | 0.869483 |
| KYY2C2 | 0.768484 | 0.905281 | 0.896817 |
| TYY2C2 | 0.768484 | 0.891124 | 0.915158 |
| KNF2C2 | 0.768484 | 0.976982 | 0.830015 |
| TNF2C2 | 0.768484 | 0.958879 | 0.844322 |
| KYF2C2 | 0.768484 | 0.930465 | 0.858645 |
| TYF2C2 | 0.768484 | 0.917383 | 0.869483 |
| KNY1Y2 | 0.414973 | 0.895737 | 0.795336 |
| TNY1Y2 | 0.414973 | 0.864359 | 0.828863 |
| KYY1Y2 | 0.414973 | 0.771709 | 0.877727 |
| TYY1Y2 | 0.414973 | 0.735268 | 0.907645 |
| KNF1Y2 | 0.414973 | 0.948359 | 0.723351 |
| TNF1Y2 | 0.414973 | 0.892625 | 0.753201 |
| KYF1Y2 | 0.414973 | 0.824725 | 0.795336 |
| TYF1Y2 | 0.414973 | 0.794297 | 0.828863 |
| KNY2Y2 | 0.414973 | 0.895737 | 0.646288 |
| TNY2Y2 | 0.414973 | 0.864359 | 0.684442 |
| KYY2Y2 | 0.414973 | 0.771709 | 0.75155 |
| TYY2Y2 | 0.414973 | 0.735268 | 0.795336 |
| KNF2Y2 | 0.414973 | 0.948359 | 0.570356 |
| TNF2Y2 | 0.414973 | 0.892625 | 0.604481 |
| KYF2Y2 | 0.414973 | 0.824725 | 0.646288 |
| TYF2Y2 | 0.414973 | 0.794297 | 0.684442 |

#### 4 Sensitivity Analyses and Outcome MeasuresScenario Evaluations

The main analysis consisted of simulating 100 realisations of 100,000 individuals living in a particular transmission setting (PfPR=1%, 5%, 10%, 20%), with varying access to antimalarial drugs if febrile (coverage=20%, 40%, 60%). One primary ACT was used as first-line therapy (DHA-PPQ, ASAQ, AL) and five different allele frequencies of pre-existing partner drug resistance (0.0, 0.01, 0.10, 0.25, 0.50) at time = 0 were explored, resulting in 18,000 simulations by each model. For our sensitivity analyses, we chose one transmission setting (PfPR = 20%) and one treatment coverage level (40%) and conducted 50 realisations of 100,000 individuals, with each sensitivity analysis scanning across seven parameter values (yielding 10,500 simulations per sensitivity analysis by each model) to characterise the impact of the following drivers of resistance.

##### 4.1 Resistant genotypes' fitness costs

The default daily fitness cost of resistance, $c$, was assumed to be equal to 0.0005. The fitness of the wild-type parasite KNY1C1 is defined as $1.0$, and the fitness of a resistant strain is defined as $1.0 - c$. In the sensitivity analysis, we explored increasing and decreasing $c$ by 10%, 20% and 30%. If a model has multiple fitness costs (for example affecting within-host parasite densities and/or onward probability of transmission) all fitness costs will be increased or reduced by the same amount.

##### 4.2 Genotype-specific drug efficacies

To explore the impact of drug efficacies, we altered our assumptions about the default efficacy of each ACT on each parasite genotype. We explored increasing and decreasing the efficacy of each drug by 15%, 10% and 5%. In the PSU model, this was modelled by varying the EC50 values by 5%, 10%, and 15%, generating 6 new drug efficacy by genotype tables. These tables were used directly by the Imperial model giving altered probabilities of 28-day treatment failure. Lastly, the MORU model varied the killing rate by 5%, 10% and 15%.

##### 4.3 Duration of infection of asymptomatic carriage

To characterise the role of asymptomatic infections, we varied the duration of asymptomatic infections in the third sensitivity analysis. Asymptomatic infections, resulting from both drug treatment failure and infections that did not result in a clinical case of infection, were increased or decreased by 10%, 20%, 30%. In the Imperial model this was included by altering the duration of asymptomatic infections, while leaving the duration of the following sub-patent infection the same. In the MORU model, the mean duration of an untreated malaria infection was altered.

##### 4.4 Probability of progressing to symptoms after an infectious bite

The rate at which new infections seek treatment will affect the selective pressure on antimalarial resistance. To explore this, we increased or decreased the probability that an individual will develop a clinical infection, i.e. an infection that is sufficiently severe in order to require treatment, by 5%, 10% and 15%. These changes alter only the probability of developing symptoms and not the probability that an individual will seek treatment, which remains fixed at 40% (treatment coverage). In all models, this change was implemented by altering both the maximum and minimum probability of developing symptoms dependent on the individual's immunity.

---

As in the main analysis, the main reported metric is the allele frequency of 580Y (a proxy for artemisinin resistance). Allele frequency was calculated as the weighted number of parasite-positive individuals carrying the 580Y allele / total number of parasite-positive individuals. The weights for each person describe the fraction of their clonal populations carrying 580Y, e.g. an individual host with five clonal infections two of which carry 580Y would be given a weight of 0.4.

#### Supplementary Figures

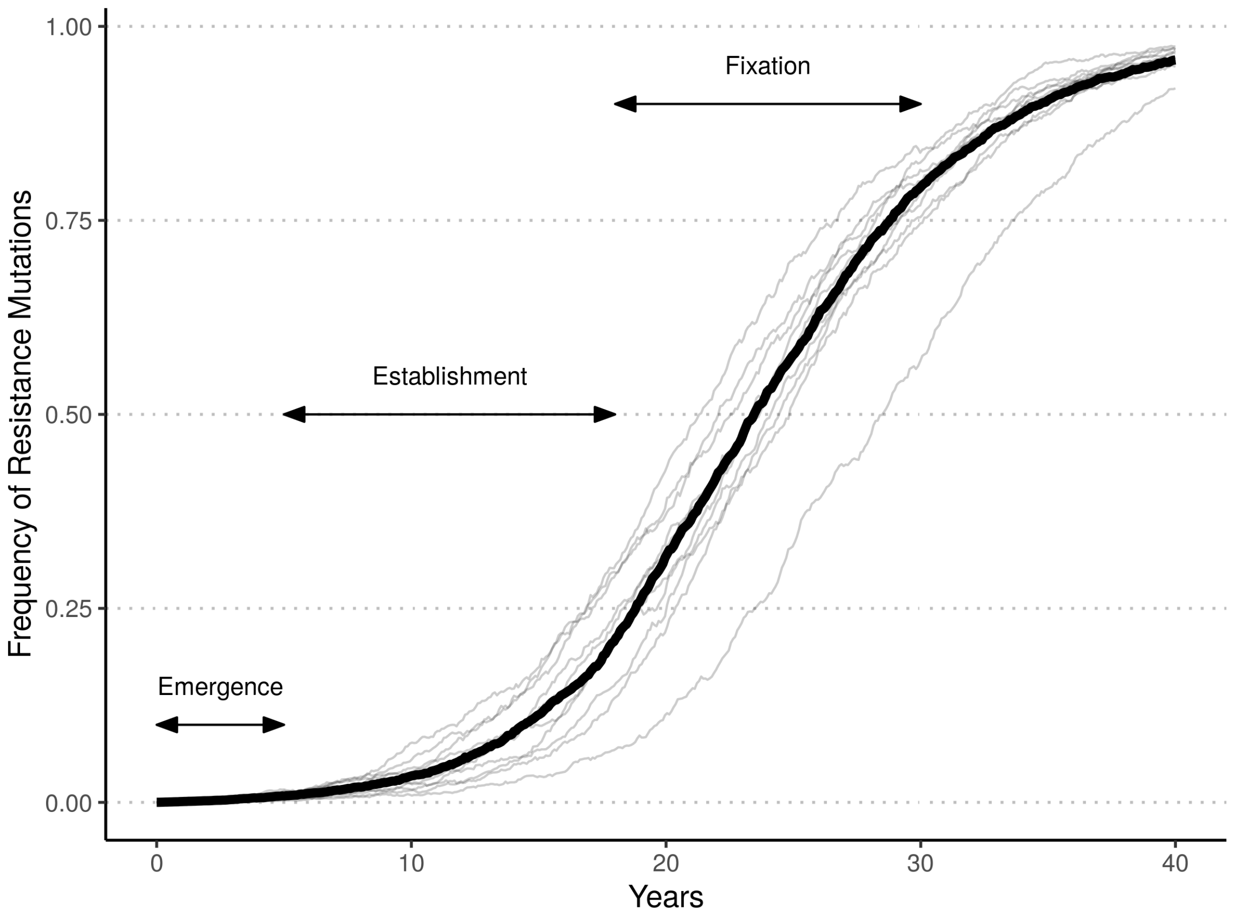


**Figure 1. Stages of drug resistance development.** Drug resistance development is often typified by three phases describing the speed at which resistance increases; Emergence, Establishment and Fixation. Prior to emergence, mutations must first arise that confer drug resistance. Due to the rarity of these events and the low effective population size of mutant strains during emergence, this stage may vary in its length due to stochastic effects. After mutant strains have persisted through a number of transmission cycles, the spread of resistance occurs leading to the rapid increase in mutation frequency during establishment, during which stochastic effects are minimised. Following establishment, the increase in resistance mutations slows until they reach fixation. An example of ten stochastic repetitions from our study for one scenario are shown with the median resistance frequency increase shown in bold. The timing of emergence (frequency 0 - 0.01), establishment (frequency 0.01 - 0.25) and fixation (frequency 0.25 - 1.00) are shown with two headed arrows.

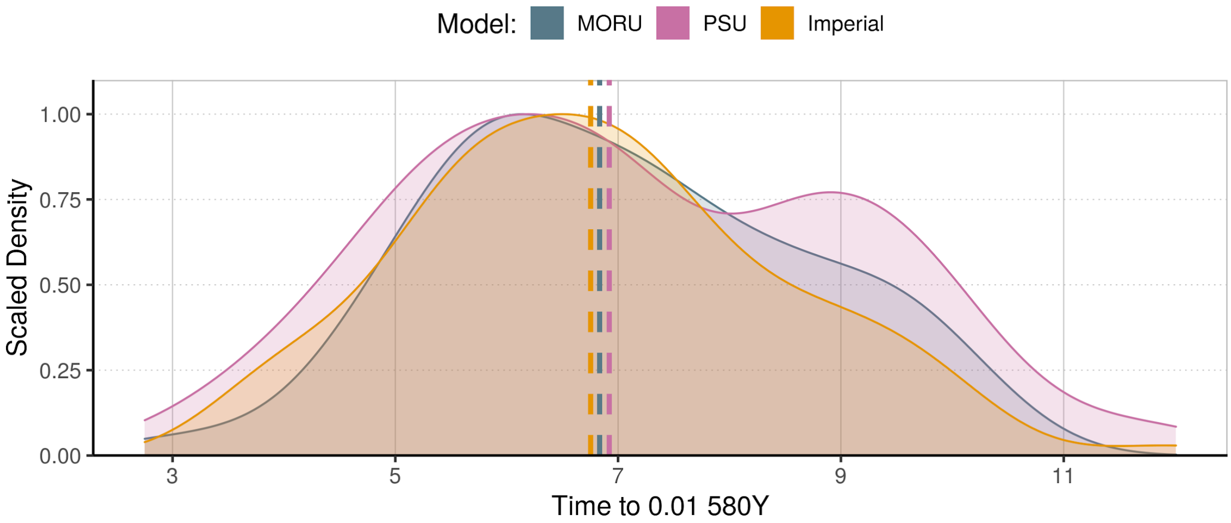


**Figure 2. Alignment exercise for 7 years till 0.01 580Y frequency.** Scaled density for model times to 0.01 frequency of 580Y as part of model alignment exercise. We aligned the three models' de novo mutation rates forcing the models to reach 0.01 allele frequency of 580Y (i.e. early artemisinin resistance emergence) after seven years exactly under a specified set of conditions: 100,000 individuals in a transmission setting with all-ages PfPR=10% and 40% coverage with DHA-PPQ as a first-line therapy.

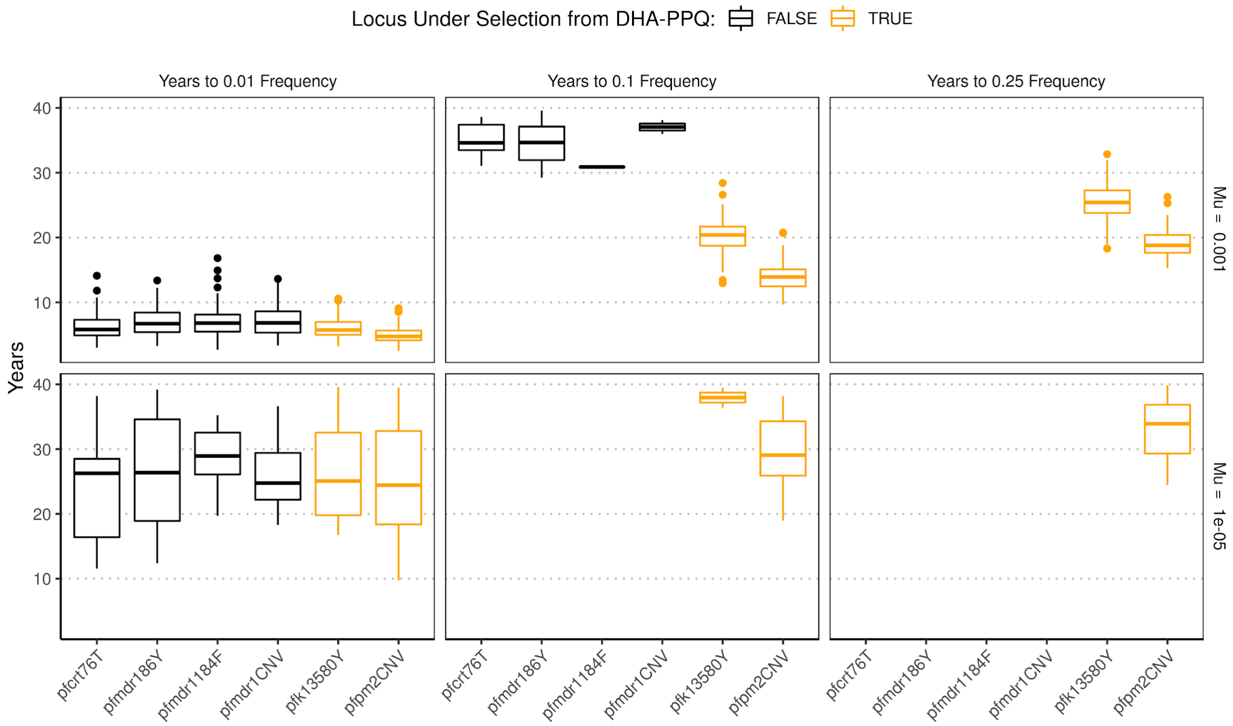


**Figure 3: Demonstration of equivalent emergence times for artemisinin resistance and loci not under selection.** Distribution of times till different drug resistance frequency milestones (0.01, 0.1, 0.25 frequency) for each locus considered in a scenario in which DHA-PPQ is the first-line therapy. All clones are the wild-type genotype at the beginning of the simulation, with the same mutation rate (shown in rows) assumed for each locus. The two loci under selection by DHA-PPQ (pfk13-580Y and pfpm2-CNV) are shown in gold and the other loci, which confer no fitness cost or resistance advantage in the scenarios considered are shown in black. An approximately equivalent time until 0.01 resistance frequency is observed for all loci, indicating that the time to emergence (0.01) is driven largely by the mutation rate chosen rather than the impacts of selection on arising mutations. For larger drug resistance milestones (0.1, 0.25) the resistance phenotype associated with pfk13-580Y and pfpm2-CNV results in shorter times till 0.1 and 0.25 frequency, indicating that these times are driven by the resistance phenotype rather than the mutation rate chosen.

**a**

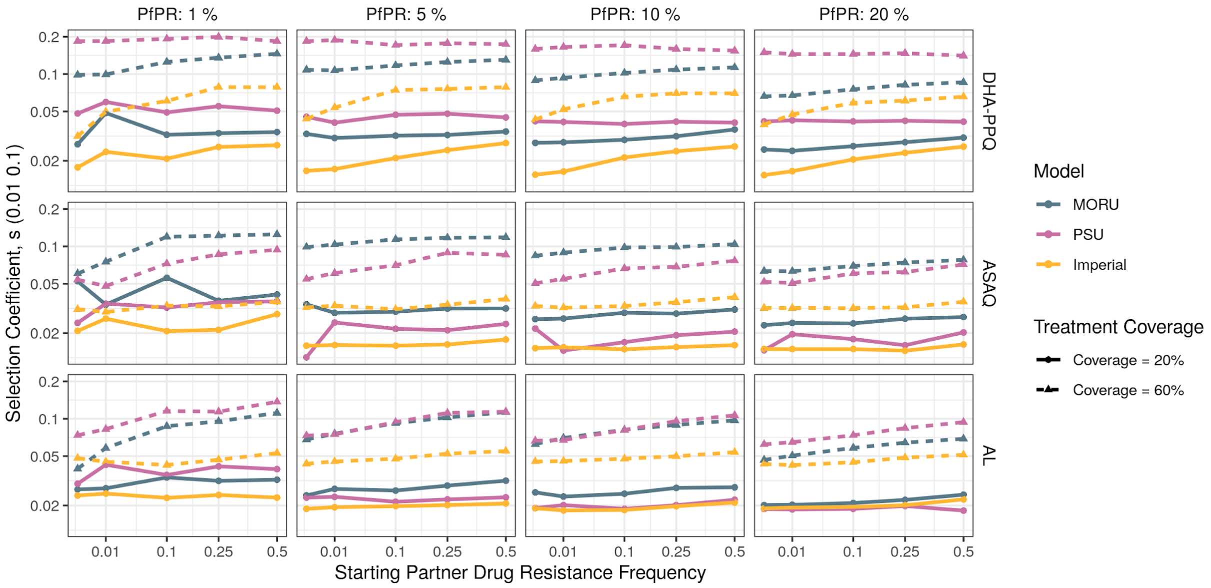


**b**

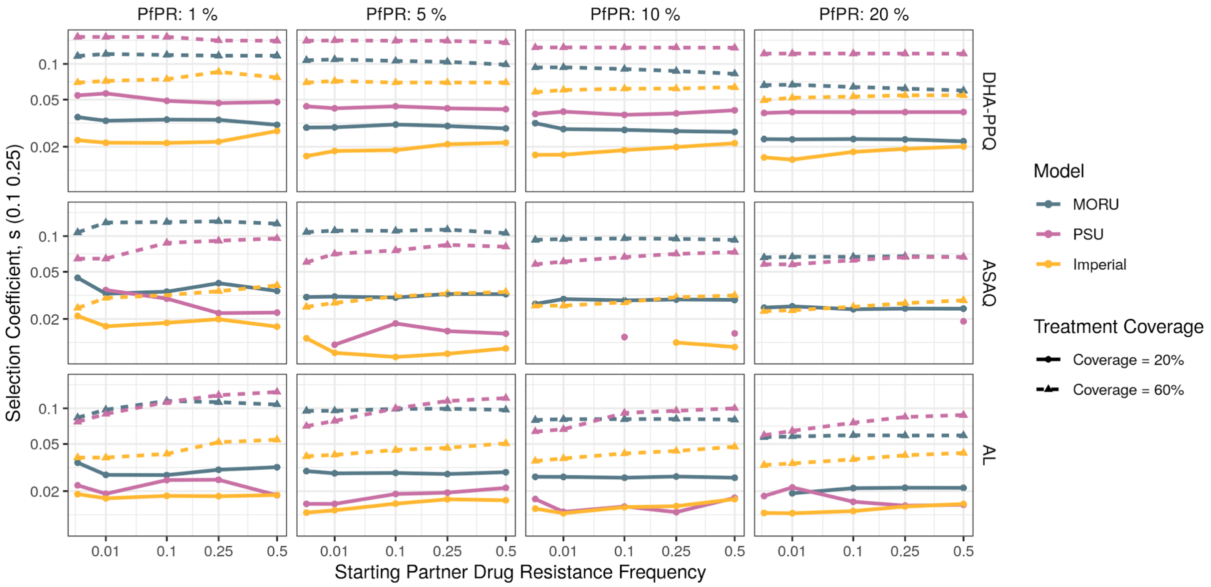


**Figure 4: Impact of starting partner drug resistance frequency on selection coefficients of 580Y**. Median selection coefficients are shown, calculated using time taken for artemisinin resistance to increase from **a)** 1% to 10% ($s_{0.01, 0.1}$) and **b)** 10% to 25% ($s_{0.1, 0.25}$). Line type and shape of point show the assumed treatment coverage and the color indicates the different transmission models used. Each column shows a different parasite prevalence and each row is a different assumed first-line ACT. For the simplest drug resistance mechanism (DHA-PPQ) the selection coefficients are notably flatter with respect to partner drug resistance in **b)** reflecting the constant rate of selection once resistance has reached the exponential stage of growth and is unaffected by stochastic fluctuations during the emergence of resistance. These dynamics are less pronounced for AL and ASAQ, which display increased selection coefficients with increasing partner drug resistance.

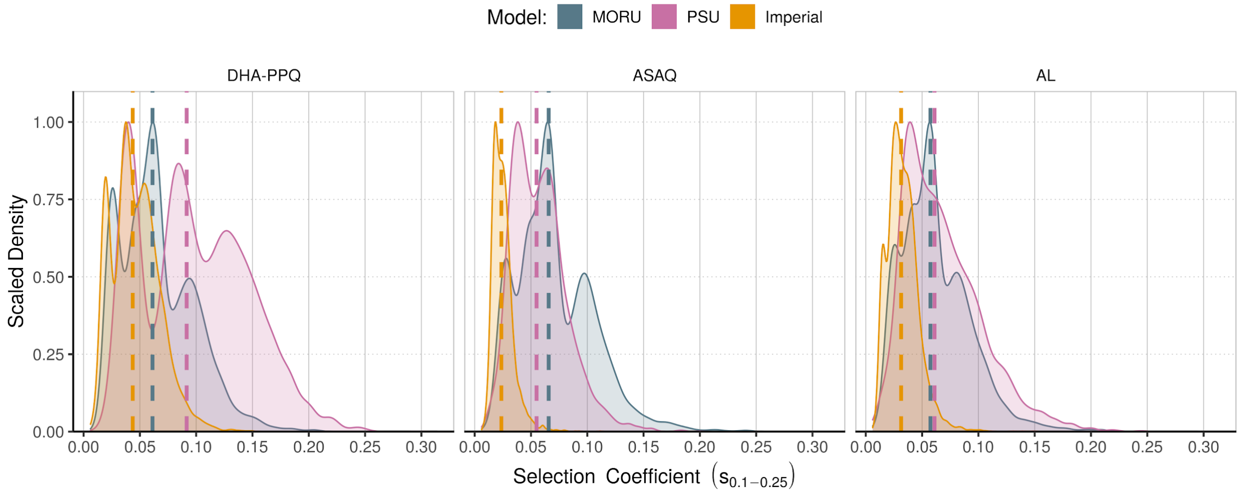


**Figure 5: Comparison between model selection coefficients.** The selection coefficients are shown for each model and first-line drug, calculated using time taken for artemisinin resistance to increase from 10% to 25% ($s_{0.1-0.25}$). The scaled density shown reflects the overall selection coefficient across all scenarios considered (malaria prevalence, starting partner drug resistance and treatment coverage), with the median selection indicated with a vertical dashed line.

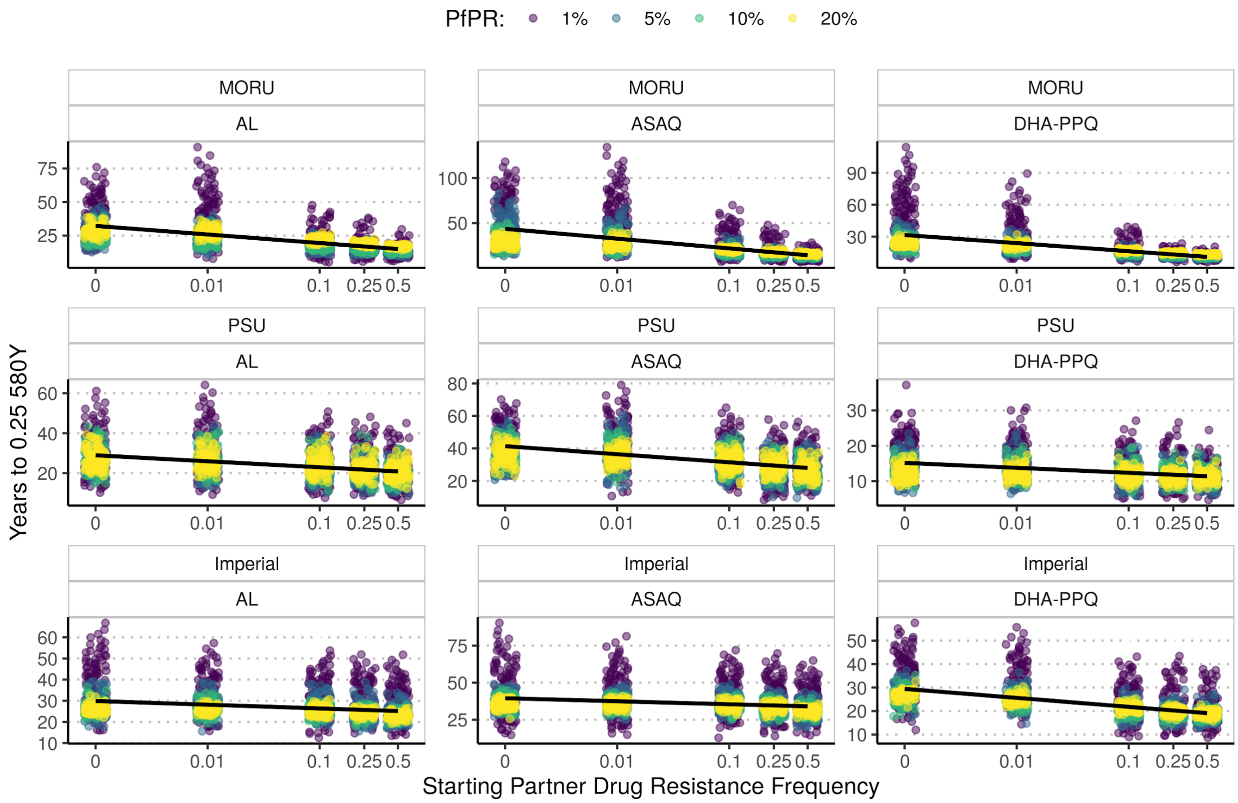


**Figure 6: Time (years) to artemisinin resistance establishment.** The time for 580Y to reach 0.25 frequency is shown for each model and first line therapy. The points shown are jittered to ease representation of the variance in times observed for each starting partner drug resistance frequency (0, 0.01, 0.1, 0.25, 0.5). The parasite prevalence (PfPR) is shown by the colour of the points revealing greater variance in emergence times at lower parasite prevalence. The approximate log linear relationship is indicated by the black line.

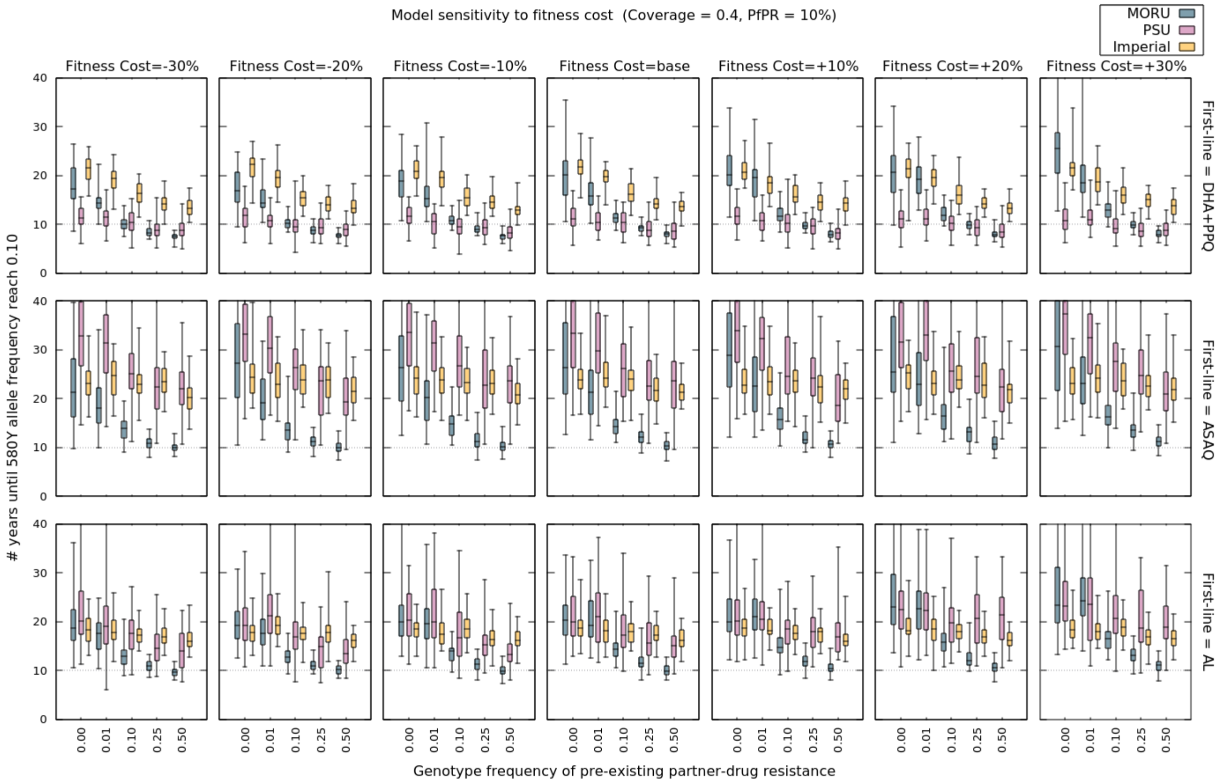


**Figure 7:** Number or years until 580Y allele frequency to increase from 0.00 to 0.25 in regions depending on assumed fitness cost of resistance.

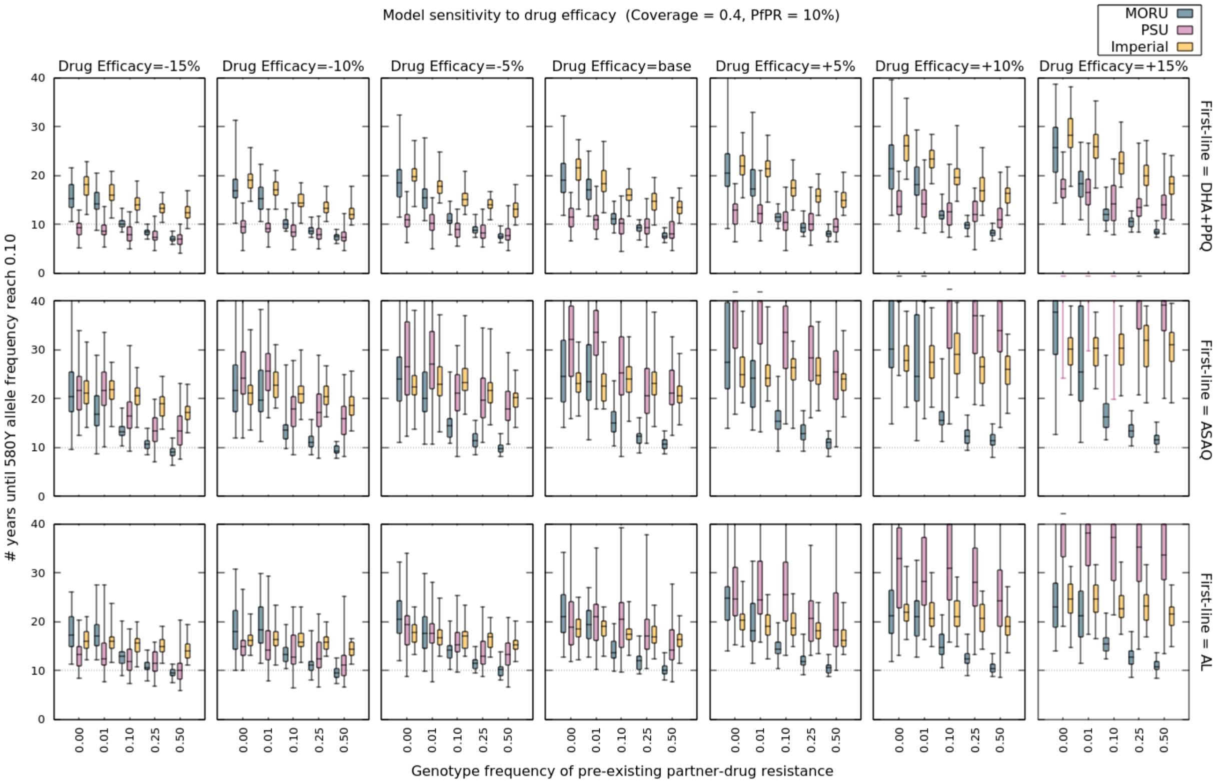


**Figure 8:** Number or years until 580Y allele frequency to increase from 0.00 to 0.25 in regions depending on assumed drug efficacy.

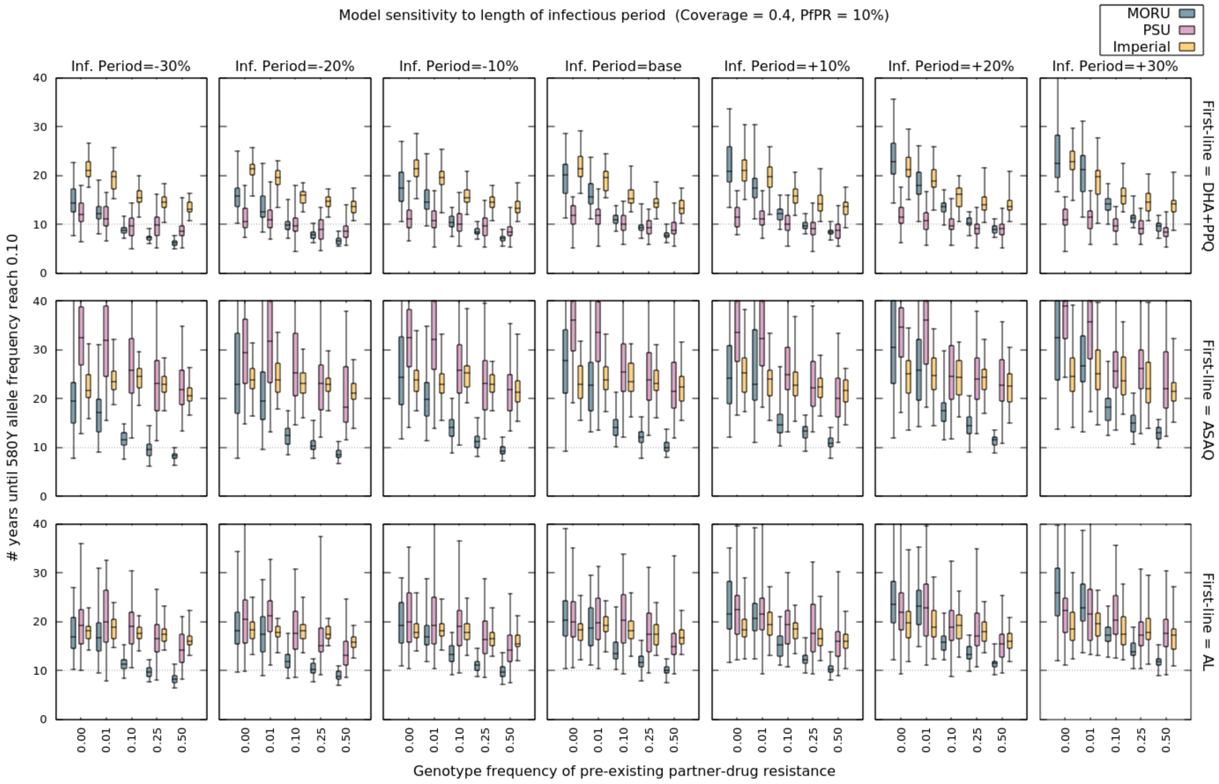


**Figure 9:** Number or years until 580Y allele frequency to increase from 0.00 to 0.25 in regions depending on assumed infectious period.

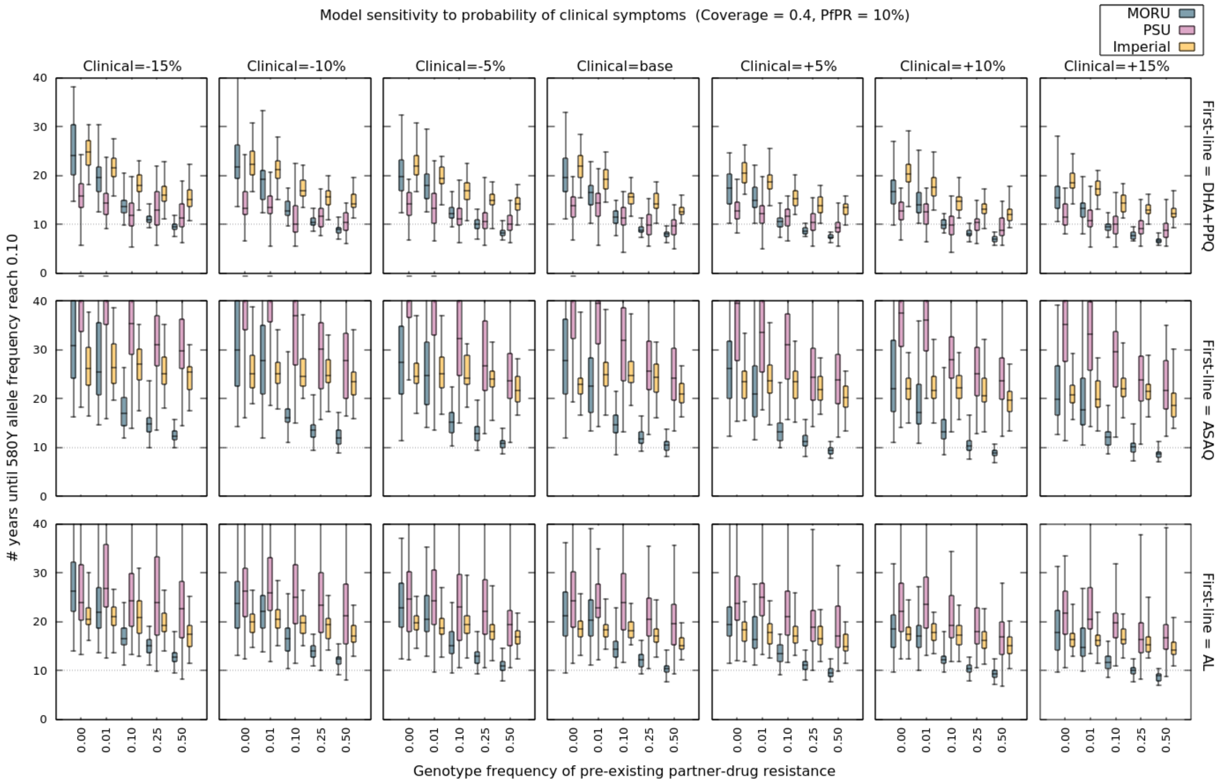


**Figure 10:** Number or years until 580Y allele frequency to increase from 0.00 to 0.25 in regions depending on assumed probability of developing clinical symptoms.

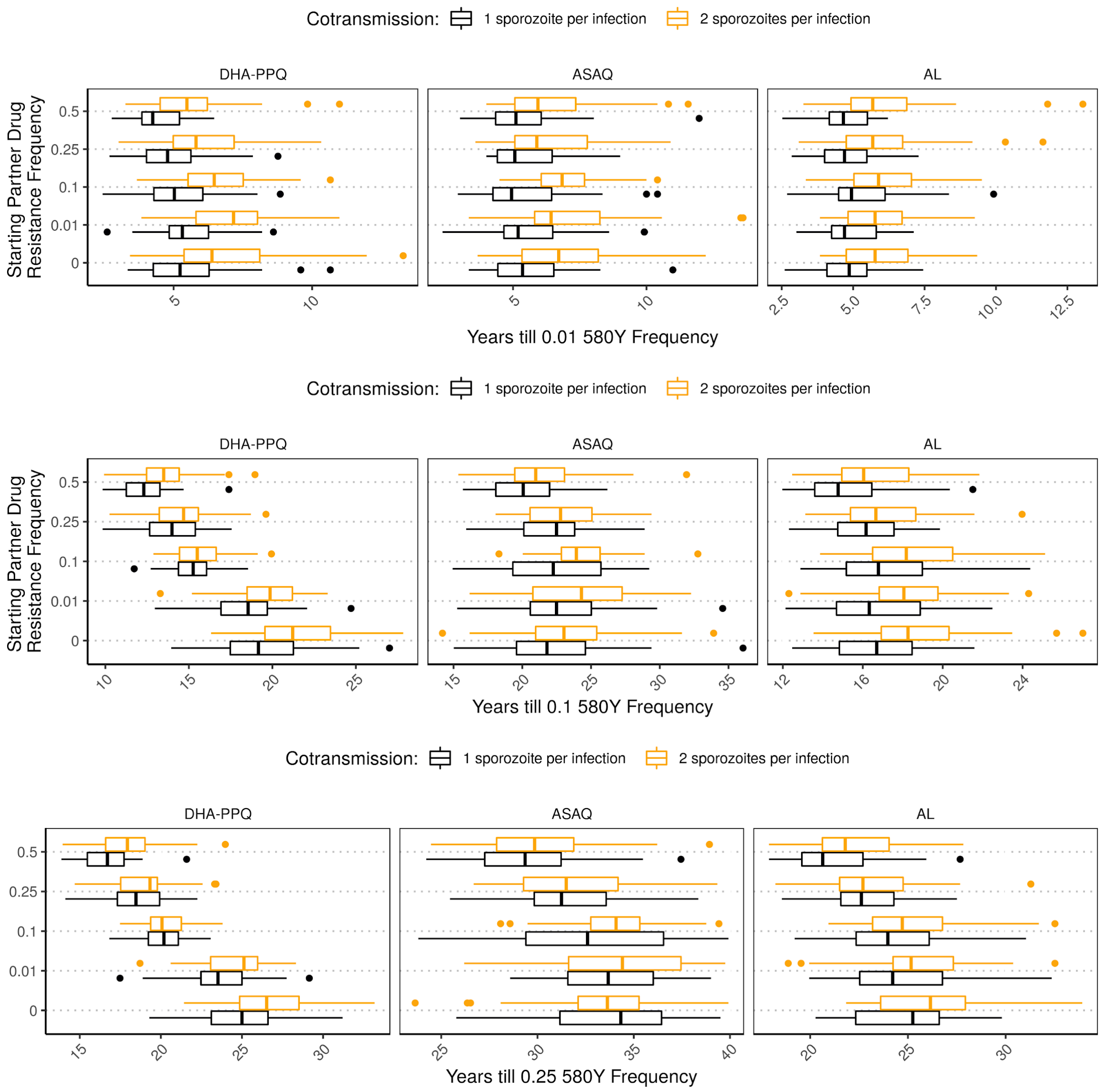


**Figure 11: Comparison of selection speed for AL and ASAQ under reduced cotransmission of sporozoites.** The effect of reducing the frequency of cotransmission events (multiple sporozoites being transmitted from one feeding event) on the speed of selection in the Imperial model is shown for each drug strategy and drug resistance milestone. Reducing cotransmission increases selection due to the reduction in interclone competition manifested by a reduction in the probability that an emergent resistant parasite will be sampled when a polyclonally infected individual is bitten by a mosquito.

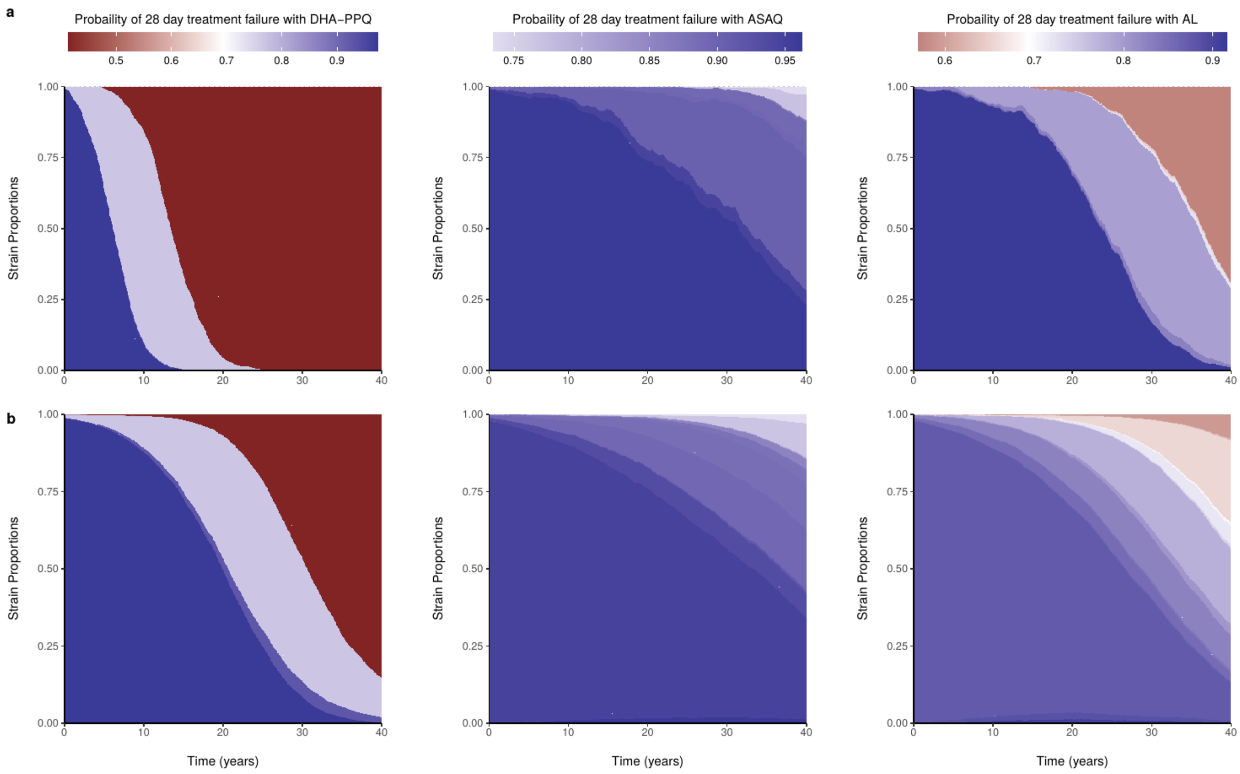


**Figure 12: Clonal interference differences dependent on mutation assumptions.** The proportion of each strain is shown over time for one stochastic repetition of **a)** the PSU model and **b)** the Imperial model. The mean probability of treatment failure after 28-days for each strain is shown, indicating the increasing treatment failure over time as wildtype parasite strains (less resistant = blue) are replaced with resistant strains (more resistant = red). For treatment with AL and ASAQ, the increased complexity of the fitness landscape resulting from increased loci associated with partner drug resistance increases clonal interference between resistant strains. The assumed absence of mutations leading to less fit clones in the PSU model results in fewer clones being generated in the ASAQ and AL scenario reducing clonal interference in comparison to the Imperial model.

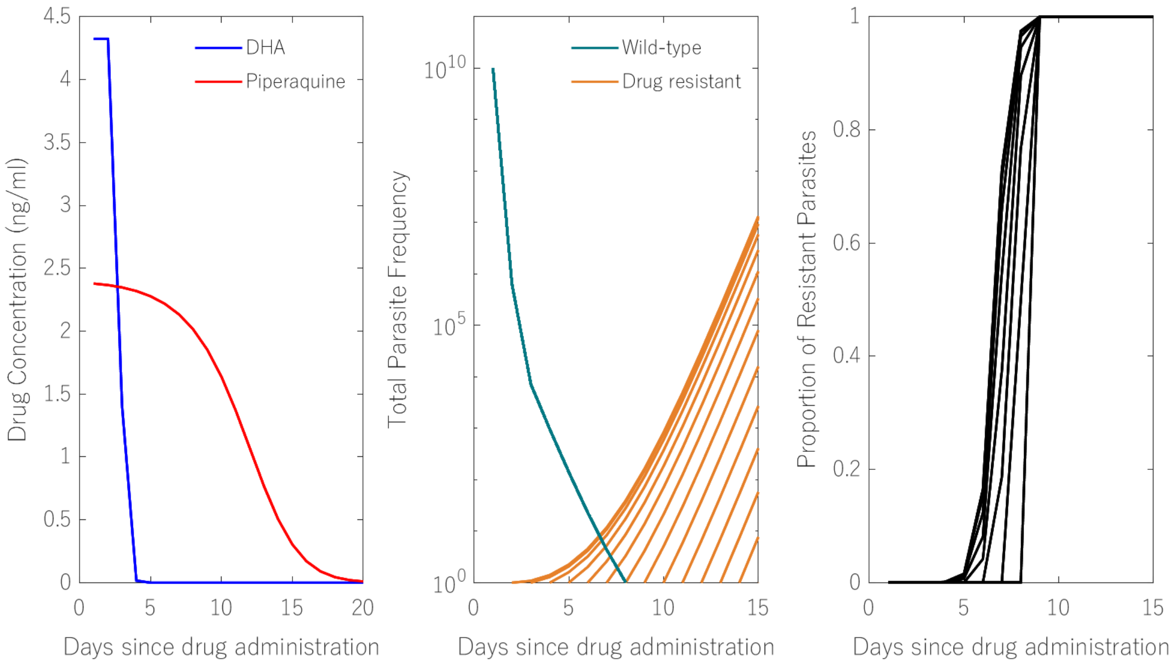


**Figure 13. Within host dynamics following a clinical infection treatment.** Here, we depict the time dynamics of drug concentration (left), and parasite population frequencies (middle, right) after treatment with DHA-PPQ. The middle panel shows how drug resistant clones (in this case, resistant to both drugs) emerging at different points in time (each line is a different simulation) will eventually outcompete wild-type parasites.

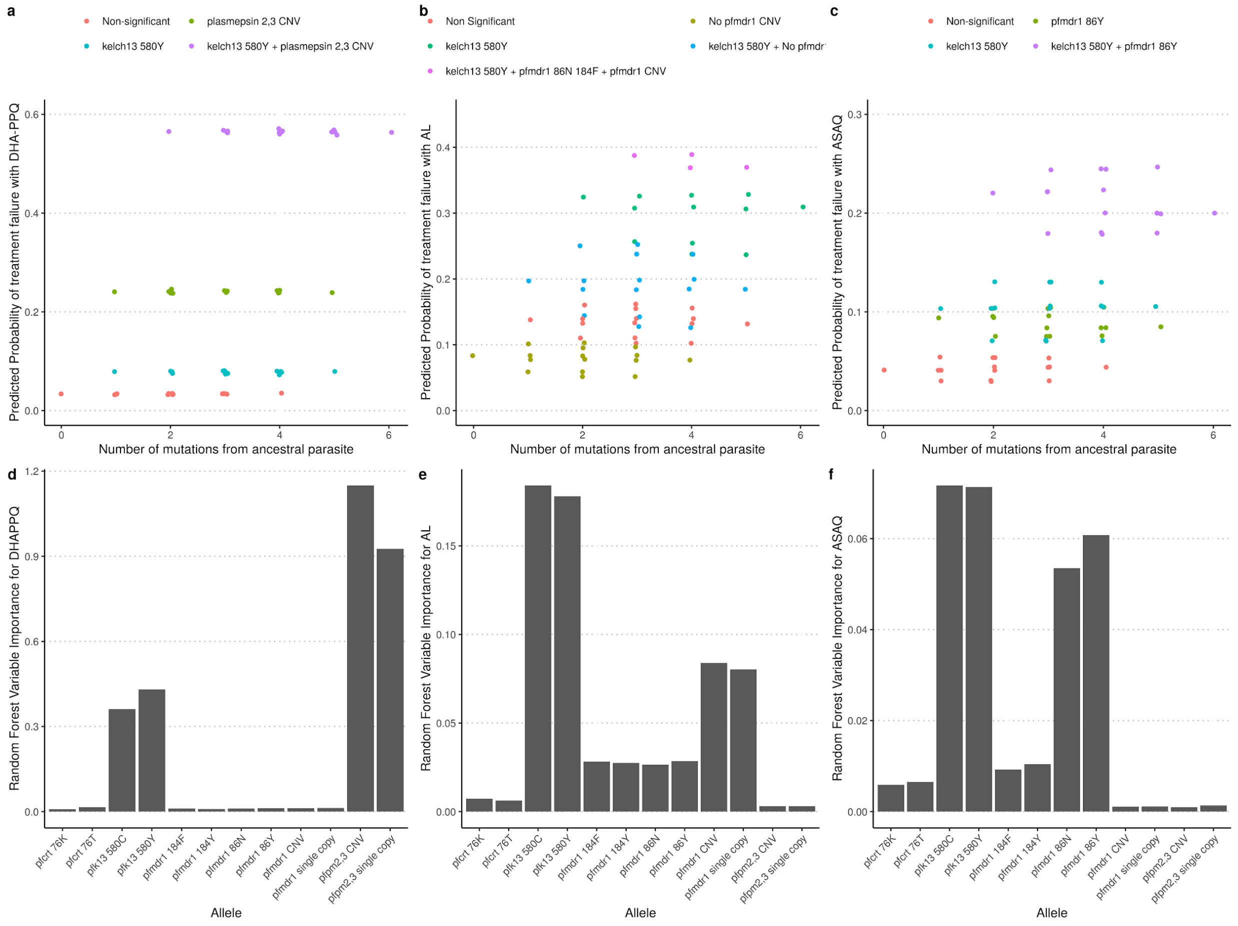


**Figure 14: Contribution of resistance alleles to treatment failure from random forest model.** The predicted treatment failure from a random forest model for **a)** DHA-PPQ, **b)** AL and **c)** ASAQ is shown for each of the 64 genotypes modelled. The model was trained using the presence/absence of each allele to predict the treatment efficacy, with the presence of specific alleles shown by the color of the points. In **d)**, **e)** and **f)** the importance of each allele towards the random forest is shown for each ACT. The variable importance is most defined for **d)** DHA-PPQ, for which only *pfkelch13* and *pfplasmepsion2,3* impact the resistance phenotype, whereas the contribution of each allele is more complex for **e)** AL and **f)** ASAQ.

#### Supplementary Tables

**Supplementary Table 4. Time till 0.25 580Y frequency ($T_{Y0.25}$) under 40% treatment coverage and 0 starting partner drug resistance.** Median and interquartile range (square brackets) is shown. Censored times above 40 years were inferred using a Weibull distribution to describe time-to-event values.

| Model | ACT | Years till 0.25 580Y frequency | | | |
|---|---|---|---|---|---|
| | | **PfPR = 1%** | **PfPR = 5%** | **PfPR = 10%** | **PfPR = 20%** |
| MORU | DHA-PPQ | 51.5 [36.7, 68.5] | 25.6 [21.4, 32.2] | 22.6 [19.4, 25.8] | 23.5 [21.8, 25.3] |
| PSU | DHA-PPQ | 19.6 [16.2, 22.3] | 13.9 [11.7, 16.1] | 13.4 [11.8, 15.5] | 12.3 [10.6, 13.5] |
| Imperial | DHA-PPQ | 36.3 [28.0, 41.9] | 27.1 [24.8, 30.7] | 26.0 [24.6, 28.2] | 27.5 [26.4, 28.2] |
| MORU | ASAQ | 64.5 [48.8, 80.1] | 41.7 [30.5, 53.3] | 28.9 [23.9, 37.7] | 27.8 [25.6, 32.0] |
| PSU | ASAQ | 46.7 [38.9, 54.0] | 38.8 [31.0, 44.0] | 38.4 [32.8, 43.2] | 36.2 [32.1, 40.6] |
| Imperial | ASAQ | 46.0 [30.1, 58.6] | 38.3 [33.2, 42.0] | 35.2 [32.5, 39.6] | 35.1 [33.3, 36.9] |
| MORU | AL | 39.8 [28.1, 49.7] | 27.2 [22.4, 34.4] | 24.5 [21.0, 28.2] | 29.4 [26.3, 31.4] |
| PSU | AL | 29.1 [20.3, 39.3] | 24.5 [21.4, 30.8] | 26.4 [23.0, 30.4] | 26.8 [24.2, 30.3] |
| Imperial | AL | 37.2 [26.4, 46.2] | 28.0 [24.1, 31.7] | 26.6 [24.9, 28.9] | 26.1 [24.9, 27.3] |

#### Supplementary References


Aguas, Ricardo, Lisa J. White, Robert W. Snow, and M. Gabriela M. Gomes. 2008. “Prospects for Malaria Eradication in Sub-Saharan Africa.” *PloS One* 3 (3): e1767.

Breiman, Leo. 2001. “Random Forests.” *Machine Learning* 45 (1): 5–32.

Bretscher, M. T., P. Dahal, J. Griffin, K. Stepniewska, Ac Ghani, and P. J. Guerin. 2019. “A Pooled Analysis of the Duration of Chemoprophylaxis against Malaria after Treatment with Artesunate-Amodiaquine and Artemether-Lumefantrine.” *medRxiv*. https://doi.org/10.1101/19002741.

Collins, W. E., and G. M. Jeffery. 1999. “A Retrospective Examination of Sporozoite- and Trophozoite-Induced Infections with Plasmodium Falciparum: Development of Parasitologic and Clinical Immunity during Primary Infection.” *The American Journal of Tropical Medicine and Hygiene* 61 (1 Suppl): 4–19.

Cook, Jackie, Nico Speybroeck, Tho Sochanta, Heng Somony, Mao Sokny, Filip Claes, Kristel Lemmens, et al. 2012. “Sero-Epidemiological Evaluation of Changes in Plasmodium Falciparum and Plasmodium Vivax Transmission Patterns over the Rainy Season in Cambodia.” *Malaria Journal* 11 (March): 86.

Cui, Liwang, Guiyun Yan, Jetsumon Sattabongkot, Yaming Cao, Bin Chen, Xiaoguang Chen, Qi Fan, et al. 2012. “Malaria in the Greater Mekong Subregion: Heterogeneity and Complexity.” *Acta Tropica* 121 (3): 227–39.

Erhart, Annette, Duc Thang Ngo, Van Ky Phan, Thi Tinh Ta, Chantal Van Overmeir, Niko Speybroeck, Valerie Obsomer, et al. 2005. “Epidemiology of Forest Malaria in Central Vietnam: A Large Scale Cross-Sectional Survey.” *Malaria Journal* 4 (December): 58.

Eyles, D. E., and M. D. Young. 1951. “The Duration of Untreated or Inadequately Treated Plasmodium Falciparum Infections in the Human Host.” *Journal. National Malaria Society* 10 (4): 327–36.

Filipe, João A. N., Eleanor M. Riley, Christopher J. Drakeley, Colin J. Sutherland, and Azra C. Ghani. 2007. “Determination of the Processes Driving the Acquisition of Immunity to Malaria Using a Mathematical Transmission Model.” *PLoS Computational Biology* 3 (12): e255.

Gao, Bo, Sompob Saralamba, Yoel Lubell, Lisa J. White, Arjen M. Dondorp, and Ricardo Aguas. 2020. “Determinants of MDA Impact and Designing MDAs towards Malaria Elimination.” *eLife* 9 (April). https://doi.org/10.7554/eLife.51773.

Griffin, Jamie T., Samir Bhatt, Marianne E. Sinka, Peter W. Gething, Michael Lynch, Edith Patouillard, Erin Shutes, et al. 2016. “Potential for Reduction of Burden and Local Elimination of Malaria by Reducing Plasmodium Falciparum Malaria Transmission: A Mathematical Modelling Study.” *The Lancet Infectious Diseases* 3099 (15): 1–8.

Griffin, Jamie T., Neil M. Ferguson, and Azra C. Ghani. 2014. “Estimates of the Changing Age-Burden of Plasmodium Falciparum Malaria Disease in Sub-Saharan Africa.” *Nature Communications* 5: 3136.

Griffin, Jamie T., T. Deirdre Hollingsworth, Lucy C. Okell, Thomas S. Churcher, Michael White, Wes Hinsley, Teun Bousema, et al. 2010. “Reducing Plasmodium Falciparum Malaria Transmission in Africa: A Model-Based Evaluation of Intervention Strategies.” *PLoS Medicine* 7 (8). https://doi.org/10.1371/journal.pmed.1000324.

Griffin, Jamie T., T. Déirdre Hollingsworth, Hugh Reyburn, Chris J. Drakeley, Eleanor M. Riley, and Azra C. Ghani. 2015. “Gradual Acquisition of Immunity to Severe Malaria with Increasing Exposure.” *Proceedings of the Royal Society B: Biological Sciences* 282:

20142657.
Gryseels, Charlotte, Lies Durnez, René Gerrets, Sambunny Uk, Sokha Suon, Srun Set, Pisen Phoeuk, et al. 2015. “Re-Imagining Malaria: Heterogeneity of Human and Mosquito Behaviour in Relation to Residual Malaria Transmission in Cambodia.” *Malaria Journal* 14 (April): 165.
Maire, Nicolas, Thomas Smith, Amanda Ross, Seth Owusu-Agyei, Klaus Dietz, and Louis Molineaux. 2006. “A Model for Natural Immunity to Asexual Blood Stages of Plasmodium Falciparum Malaria in Endemic Areas.” *The American Journal of Tropical Medicine and Hygiene* 75 (2 Suppl): 19–31.
Marsh, K., and R. Snow. 1999. “Malaria Transmission and Morbidity.” *Parassitologia* 41: 241–46.
Molineaux, L., and G. Gramiccia. 1980. “The Garki Project. Research on the Epidemiology and Control of Malaria in the Sudan Savanna of West Africa.” https://doi.org/10.1016/0035-9203(81)90085-7.
Nguyen, Tran Dang, Piero Olliaro, Arjen M. Dondorp, J. Kevin Baird, Ha Minh Lam, Jeremy Farrar, Guy E. Thwaites, Nicholas J. White, and Maciej F. Boni. 2015. “Optimum Population-Level Use of Artemisinin Combination Therapies: A Modelling Study.” *The Lancet Global Health* 3 (12): e758–66.
Nguyen, Tran Dang, Thu Nguyen-Anh Tran, Daniel M. Parker, Nicholas J. White, and Maciej F. Boni. 2021. “Antimalarial Mass Drug Administration in Large Populations and the Evolution of Drug Resistance.” *bioRxiv*. https://doi.org/10.1101/2021.03.08.434496.
Okell, Lucy C., Matthew Cairns, Jamie T. Griffin, Neil M. Ferguson, Joel Tarning, George Jagoe, Pierre Hugo, et al. 2014. “Contrasting Benefits of Different Artemisinin Combination Therapies as First-Line Malaria Treatments Using Model-Based Cost-Effectiveness Analysis.” *Nature Communications* 5: 5606.
Owusu-Agyei, S., T. Smith, H-P Beck, L. Amenga-Etego, and I. Felger. 2002. “Molecular Epidemiology of Plasmodium Falciparum Infections among Asymptomatic Inhabitants of a Holoendemic Malarious Area in Northern Ghana.” *Tropical Medicine & International Health: TM & IH* 7 (5): 421–28.
Reyburn, Hugh, Redempta Mbatia, Chris Drakeley, Jane Bruce, Ilona Carneiro, Raimos Olomi, Jonathan Cox, et al. 2005. “Association of Transmission Intensity and Age with Clinical Manifestations and Case Fatality of Severe Plasmodium Falciparum Malaria.” *JAMA: The Journal of the American Medical Association* 293 (12): 1461–70.
Ross, Amanda, Gerry Killeen, and Thomas Smith. 2006. “Relationships between Host Infectivity to Mosquitoes and Asexual Parasite Density in Plasmodium Falciparum.” *The American Journal of Tropical Medicine and Hygiene* 75 (2 Suppl): 32–37.
Rowe, A., S. Y. Rowe, R. W. Snow, E. L. Korenromp, J. R. Armstrong-Schellenberg, C. Stein, B. L. Nahlen, J. Bryce, R. E. Black, and R. W. Steketee. 2006. “The Burden of Malaria Mortality among African Children in the Year 2000.” *International Journal of Epidemiology* 35 (3): 691–704.
Saralamba, S., W. Pan-Ngum, R. J. Maude, S. J. Lee, J. Tarning, N. Lindegardh, K. Chotivanich, et al. 2011. “Intrahost Modeling of Artemisinin Resistance in Plasmodium Falciparum.” *Proceedings of the National Academy of Sciences of the United States of America* 108 (1): 397–402.
Slater, Hannah C., Jamie T. Griffin, Azra C. Ghani, and Lucy C. Okell. 2016. “Assessing the Potential Impact of Artemisinin and Partner Drug Resistance in Sub - Saharan Africa.” *Malaria Journal*, 1–11.
Smith, D. L., J. Dushoff, R. W. Snow, and S. I. Hay. 2005. “The Entomological Inoculation Rate

and Plasmodium Falciparum Infection in African Children.” *Nature* 438 (7067): 492–95.
Watson, O. J., Robert Verity, and Rich FitzJohn. 2021. *OJWatson/magenta: Release Magenta 1.3.0*. https://doi.org/10.5281/zenodo.4620137.
Watson, Oliver J., Lucy C. Okell, Joel Hellewell, Hannah C. Slater, H. Juliette T. Unwin, Irene Omedo, Philip Bejon, et al. 2021. “Evaluating the Performance of Malaria Genetics for Inferring Changes in Transmission Intensity Using Transmission Modeling.” *Molecular Biology and Evolution* 38 (1): 274–89.
